# Arc mediates intercellular synaptic plasticity via IRSp53-dependent extracellular vesicle biogenesis

**DOI:** 10.1101/2024.01.30.578027

**Authors:** Kaelan Sullivan, Alicia Ravens, Eric de Hoog, Alicia Walker, Tessa R. Kilberg, Sevnur Kömürlü Keceli, Jenifer Einstein, Mitali Tyagi, Tom Kicmal, Michael P. Hantak, Tate Shepherd, Alyson Stewart, Kenneth Lyon, Adarsh Dharan, Balázs Fábián, Monika Fuxreiter, Thomas Gallagher, Edward M. Campbell, Jason D. Shepherd

## Abstract

Current models of learning and memory have focused on cell-autonomous regulation of synaptic strength; however, intercellular signaling between cells in the brain is important for normal cognition. The immediate early gene *Arc* is a repurposed retrotransposon critical for long-term forms of synaptic plasticity and memory. Arc protein forms virus-like capsids released in extracellular vesicles (EVs) that mediate intercellular signaling of unknown function. Here, we find that long-term potentiation stimuli induce the biogenesis of Arc EVs by recruiting the I-BAR protein IRSp53, which facilitates Arc capsid assembly, trafficking, and release from actin-rich filopodial structures in dendrites. Arc EVs transfer Arc protein and mRNA to neighboring dendrites, where translation of transferred *Arc* mRNA induces a loss of surface AMPA-type glutamate receptors. These results show that Arc EVs mediate an intercellular form of synaptic plasticity that may be critical for memory consolidation and reveals a new neuronal EV biogenesis pathway.

## Introduction

Learning and experience induce the expression of a specific set of genes that implement long-lasting changes in neuronal connectivity and synapse strength^1^. The immediate-early gene *Arc* is a critical mediator of long-lasting synaptic plasticity and memory^2,3^. Arc regulates the endocytosis of AMPA-type glutamate receptors through interactions with endocytic proteins to mediate long-term depression (LTD)^4–6^ and homeostatic plasticity^7,8^. However, the expression of Arc is also induced by long-term potentiation (LTP) stimuli^9^, despite not being strictly required for LTP expression^10^. Recent studies showed that *Arc* evolved from an ancient retrotransposon and that Arc protein contains conserved structural sequences of the retroviral Gag protein^11–14^. Similar to retroviruses, Arc forms capsids that are released from neurons with an enveloped membrane in extracellular vesicles (EVs), which transfer RNAs and proteins to recipient neurons^13^. EVs are emerging as important mediators of intercellular signaling in the nervous system^15^. However, the precise mechanisms of Arc EV release and biogenesis remain unknown. Arc EVs are also critical for the transmission of pathological tau in Alzheimer’s disease^16^. Thus, a mechanistic understanding of Arc EV biogenesis and normal function will also provide insight into the processes underlying neurodegeneration.

Arc interacts with synaptic proteins, such as PSD-95 and TARP-γ2, through a common binding motif in the N-lobe^14^. Another synaptic protein, IRSp53^17^ (also known as BAIAP2), was predicted to interact with Arc in the same region^14^. IRSp53 is an I-BAR protein that induces outward membrane curvature through interactions with the actin machinery via its SH3 domain^18^. IRSp53 promotes dendritic spine formation^19^, plays a role in memory formation^20^, and is associated with psychiatric disorders such as autism and schizophrenia^21^. IRSp53 may also control EV biogenesis by regulating the release of a specific subset of EVs from the plasma membrane in non-neuronal cells^22^. Intriguingly, IRSp53 has also been implicated in HIV-1 capsid release from the plasma membrane^23,24^. Based on these data, we investigated the role of IRSp53 in Arc EV biogenesis and whether Arc EVs mediate intercellular synaptic plasticity. Cellular models of synaptic plasticity, such as LTP and LTD have focused on cell-autonomous mechanisms^25^. Here, we describe a new mode of intercellular synaptic plasticity that may underlie long-term forms of memory.

## Results

### IRSp53 directly binds Arc capsids

IRSp53 is highly expressed in neurons^26,27^ and is implicated in the assembly and release of HIV-1 capsids^23^. Since Arc and HIV-1 share structural homology^14^, we hypothesized that IRSp53 regulates Arc capsid assembly and release. To test if Arc and IRSp53 interact in the brain *in vivo*, we immunoprecipitated (IP) Arc from homogenized brain tissue in 1-month-old mice (Figure 1A) that had experienced six hours of home-cage enrichment to increase Arc expression^28,29^ (Figure S1A). IRSp53 co-IPs with Arc in brain cortical lysates (Figure 1A). Arc also co-IPs with PSD-95, a known interacting partner^30^. To investigate whether this interaction occurs through direct binding, we purified GST-tagged IRSp53 and Arc protein. GST-IRSp53 was incubated with purified Arc protein bound on a column. While Arc and IRSp53 eluted together off the column, Arc was not found in the eluted fraction when incubated with GST alone, demonstrating a direct and specific protein-protein interaction (Figure 1B).

**Figure 1.**
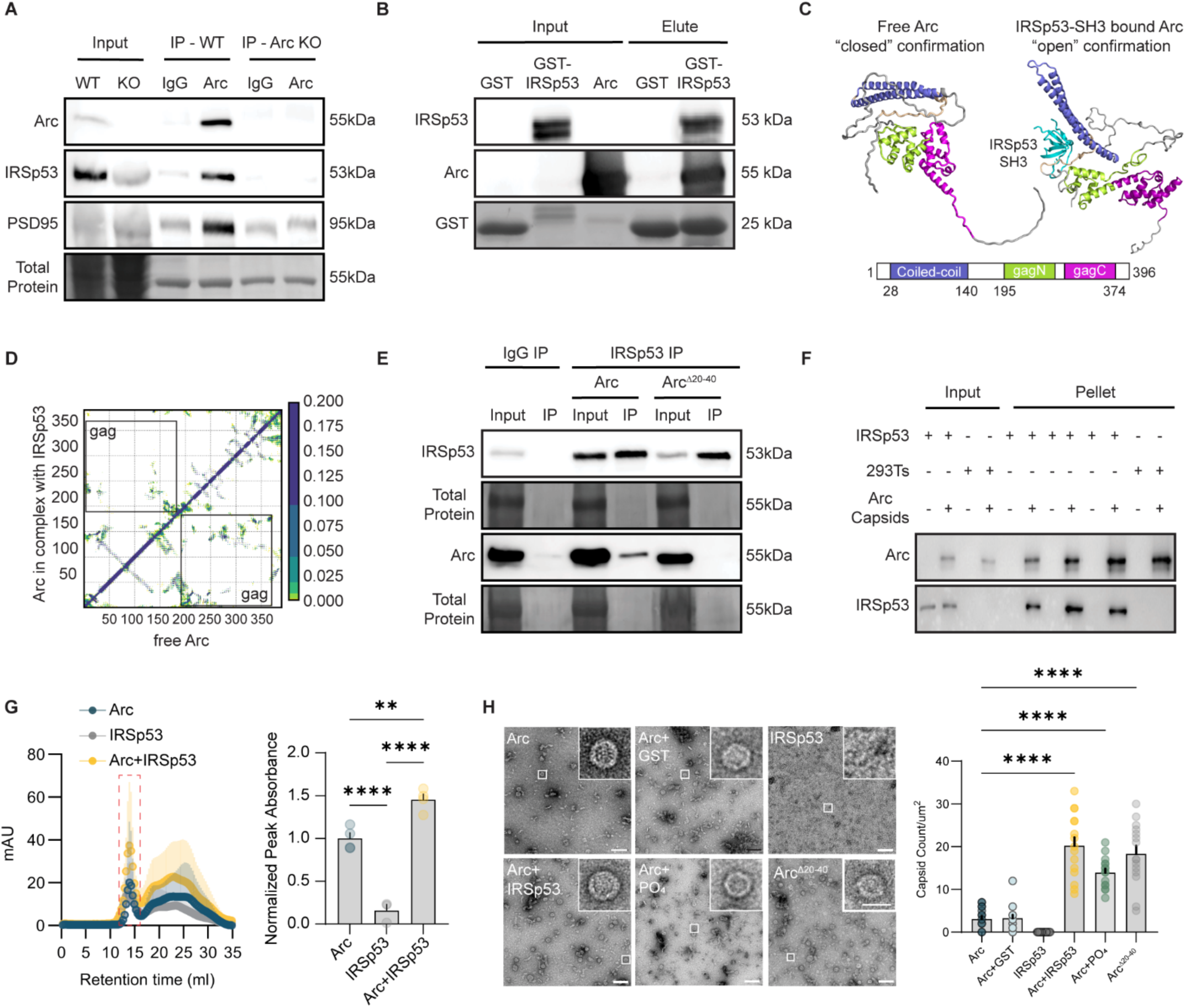
IRSp53 facilitates Arc capsid assembly via a direct protein interaction. **A**. *Arc co-IPs with IRSp53 and PSD-95*. Forebrain tissue was collected from 1-month-old WT and Arc KO mice that experienced 6 hours of environmental enrichment to increase Arc expression. Cell lysates were incubated with IgG or Arc antibody. **B**. *Arc directly binds IRSp53*. Purified Arc protein was incubated with purified GST-IRSp53 or GST (inputs) and then bound to a column. Arc eluted off the column with IRSp53 but not GST. **C**. *IRSp53 facilitates an “open” conformation when bound to Arc monomers.* Molecular dynamics model of the intramolecular interactions between the free Arc N-terminal disordered region and the N-lobe of the Gag domain (“closed” conformation) and the interactions when IRSp53 is bound that expose capsid-forming residues (“open confirmation”). **D**. *IRSp53 preferentially interacts with Arc via its SH3 domain*. Arc intramolecular interactions (< 5Å) in free Arc (*bottom triangle*) and in complex with IRSp53 (*top triangle*). In free monomeric Arc the Gag domain (grey boxes) forms contacts with the disordered N-terminal region, which are absent in the presence of IRSp53. **E**. *The Arc N-terminal coiled-coiled domain is required for IRSp53 binding.* HEK293T cells were transfected with IRSp53 and either myc-Arc or myc-Arc^Δ20–40^. Cell lysates were IPed with an IgG or IRSp53 antibody. Arc, but not Arc^Δ20–40^ co-IPed with IRSp53. **F**. *IRSp53 binds Arc capsids.* Whole-cell lysates from HEK293T cells overexpressing IRSp53 were incubated with purified Arc capsids, and the mixture was pelleted by ultracentrifugation. IRSp53 protein was only found in the pellet when Arc capsids were present. **G**. *IRSp53 increases Arc higher molecular weight species.* Chromatogram showing the peaks of purified proteins run through the size exclusion column that were selected for TEM (red box). Peak height was normalized to protein input for each experiment (Arc and IRSp53 n=4, GST n=3). **H**. *IRSp53 facilitates Arc capsid assembly.* Representative TEM images of purified proteins (0.5mg/ml concentration): Arc, Arc+GST, IRSp53, Arc+IRSp53 (1:1), Arc^Δ20–40^, all in TBS buffer, and Arc in 500mM phosphate buffer. IRSp53 enhances Arc capsid assembly (n=15 images/grid). Data are the mean ± s.e.m. One-way analysis of variance (ANOVA) with Tukey’s multiple comparisons test (**G, H**) \*\**P*>0.001, \*\*\*\**P*<0.0001, ns=not significant.

**Supplementary Figure 1.**
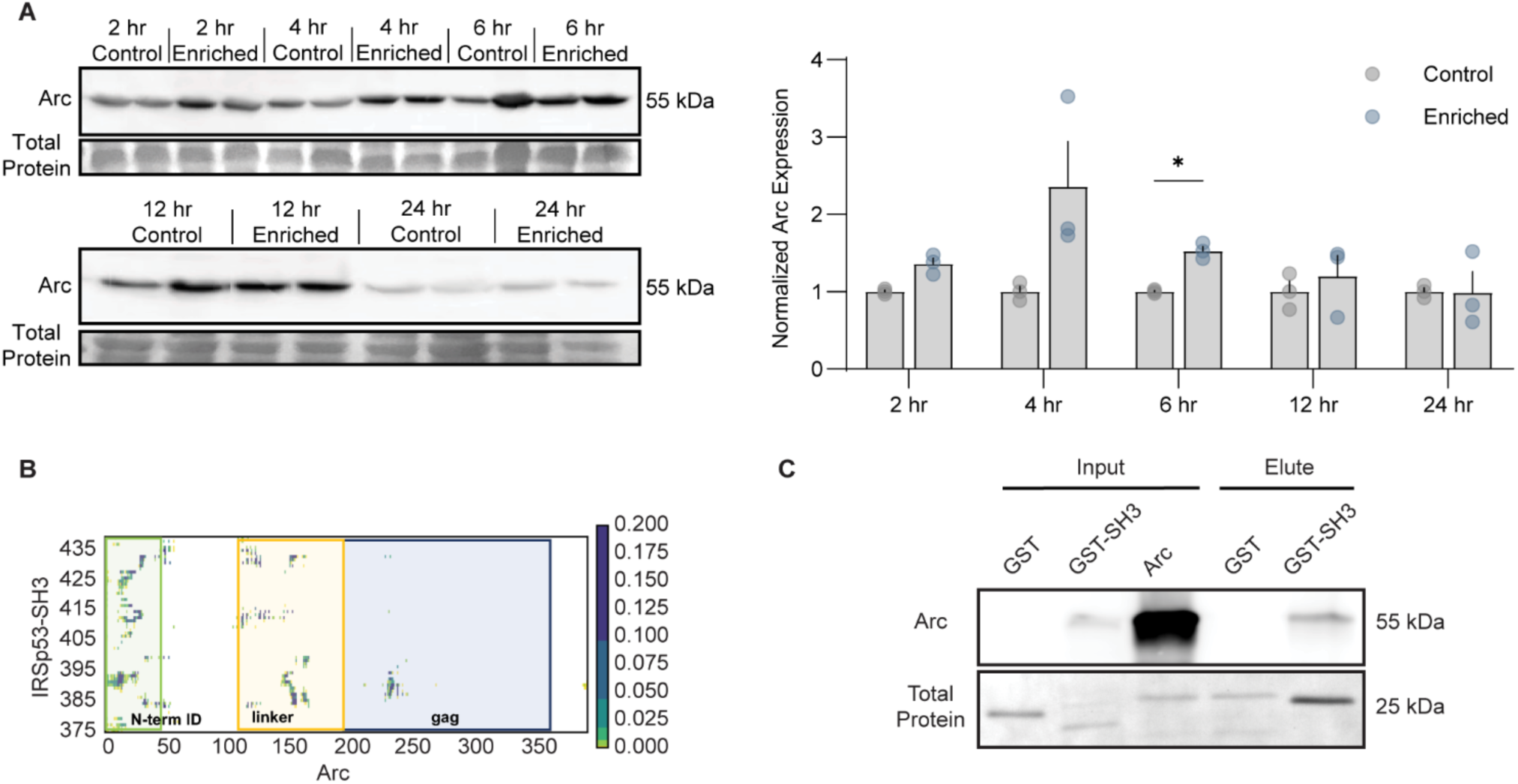
Validation of Arc and IRSp53 interactions. **A**. *Arc levels are increased with 4- and 6-hours of enriched environment.* Toys were placed in the home cage of P30 WT mice at the start of their active cycle. Enrichment toys were left in the cage until the animals were sacrificed. Forebrain tissue was collected at 2-, 4-, 6-, 12-, and 24-hours post-enrichment or from unenriched controls (n=3 per group/per timepoint). Arc bands were first normalized to total protein and then to their respective timepoint control. **B**. *The Arc N-terminal region contains various proline-rich peptide motifs that bind the SH3 domain.* Contact maps derived from the full-length IRSp53 and Arc molecular dynamics simulations. Contacts between heavy atoms < 5Å are indicated, and the fraction of contacts is shown by the scale bar (right). The IRSp53 SH3 domain contacts the N-terminal intrinsically disordered (ID) region (green) of Arc and the disordered linker (yellow), connecting the coiled-coil and Gag domains (blue). **C.** *Arc binds the IRSp53 SH3 domains.* Purified full-length Arc protein was incubated with purified GST-IRSp53-SH3 and bound to a column. Arc was eluted off the column with IRSp53-SH3 but not with GST. Data are the mean ± s.e.m. Two-way ANOVA with Šídák’s multiple comparison test (**A**) was performed. \**P*<0.05.

To gain molecular insights into Arc-IRSp53 interactions, we conducted all-atom molecular dynamics (MD) simulations using AlphaFold3^31^ predicted structural models of mouse Arc (UniProt: Q9WV31) and IRSp53 (UniProt: Q8BKX1) sequences. We generated a structural ensemble of 500 models for free monomeric Arc and in complex with IRSp53 (Figure 1C). Simulations of free Arc showed an intramolecular interaction between the coiled-coil (CC) and Gag domains (Figure 1C), mediated by the disordered Arc N-terminal region and disordered linker residues preceding the Gag domain (Figure 1D). Thus, free monomeric Arc is normally in a “closed” conformation, where the CC domain shields key Gag residues important for capsid assembly. Both structure predictions and MD simulations indicate that IRSp53 preferentially interacts with Arc via its SH3 domain, potentially through multiple sites (Figure 1D). The Arc N-terminal region contains various proline-rich peptide motifs that bind the SH3 domain during the simulations of the complex (Figure S1B). Binding of IRSp53 diminishes the intramolecular interactions between the CC and Gag domains (Figure 1D), which results in an “open” conformation where the Gag residues are exposed to potentially promote capsid assembly (Figure 1C). These data predict that IRSp53 induces a switch from the “closed” autoinhibitory state of free monomeric Arc into an “open” capsid-prone structure. Consistent with the modeling, we found that purified Arc protein binds the purified IRSp53 SH3 domain (Figure S1C). As predicted by the model, an N-terminal Arc deletion mutant (Arc ^Δ20–40^) was unable to bind to IRSp53, showing the necessity of these regions for this interaction (Figure 1E). To test whether IRSp53 interacts with the capsid form of Arc, we incubated recombinant purified Arc capsids with HEK293T cell lysates from cells transfected with IRSp53 and then the capsids were pelleted using ultracentrifugation. IRSp53 was only found in the pellet when Arc capsids were present (Figure 1F). These results show that Arc and IRSp53 interact in the brain *in vivo* and that this interaction occurs through direct binding to Arc capsids.

### IRSp53 facilitates Arc capsid assembly

Since IRSp53 binds Arc capsids, we determined whether IRSp53 facilitates capsid assembly. We incubated purified Arc and IRSp53 in a 1:1 molar ratio in Tris-buffered saline solution (TBS), a buffer not conducive for capsid assembly^13^. We then performed size exclusion chromatography (SEC) and collected the peak protein fractions (Figure 1G, red box). In the chromatogram of eluted proteins, we observed an increase in the void fraction peak height of IRSp53-incubated Arc samples, consistent with a shift from low- to high-order Arc oligomers (Figure 1G). The void fractions were then processed for negative stain transmission electron microscopy (TEM). Arc protein alone or when incubated with GST in TBS rarely formed capsids, and no capsids were seen on grids with only IRSp53 (Figure 1H). In contrast, when IRSp53 was incubated with Arc we observed a significant increase in Arc capsid formation in TBS, similar to Arc in high phosphate buffer (Figure 1H). To test the prediction that the Arc N-terminal CC domain maintains monomeric Arc in a “closed” state, we purified Arc^Δ20–40^ and performed TEM. Arc^Δ20–40^ readily forms capsids in TBS buffer at similar levels to when IRSp53 is added to full-length Arc, suggesting disinhibition of the autoinhibitory state via CC and Gag interactions (Figure 1H). These data indicate that IRSp53 acts as a molecular switch, facilitating capsid assembly by releasing monomeric Arc’s autoinhibitory state into an “open” capsid prone state.

### LTP induces IRSp53-dependent anterograde Arc trafficking and release from dendrites

We next determined where and when Arc interacts with IRSp53 in neurons. We used a proximity ligation assay (PLA) to detect endogenous protein-protein interactions between Arc and IRSp53 in primary rat hippocampal neurons (Figure 2A). Since Arc and IRSp53 expression in neurons is regulated by neuronal activity^9,26^, we treated neurons with chemical induction of LTP or LTD (cLTP/cLTD)^32,33^. Arc-IRSp53 puncta increased after cLTP, but not after cLTD (Figure 2A). Arc expression increases in dendrites after both cLTP and cLTD (Figure S2A-B), whereas IRSp53 expression and IRSp53-Arc colocalization only increased in dendrites after cLTP (Figure S2A-D). Arc and IRSp53 colocalize specifically in actin-rich extra-synaptic areas in dendrites after cLTP that are not present in basal or cLTD conditions (Figure S2E, G). In contrast, Arc colocalization increases at synapses after cLTD (Figure S2F, H). These data show that LTP stimuli preferentially induce Arc and IRSp53 interactions in the cell body and in actin-rich extra-synaptic sites in dendrites.

**Figure 2.**
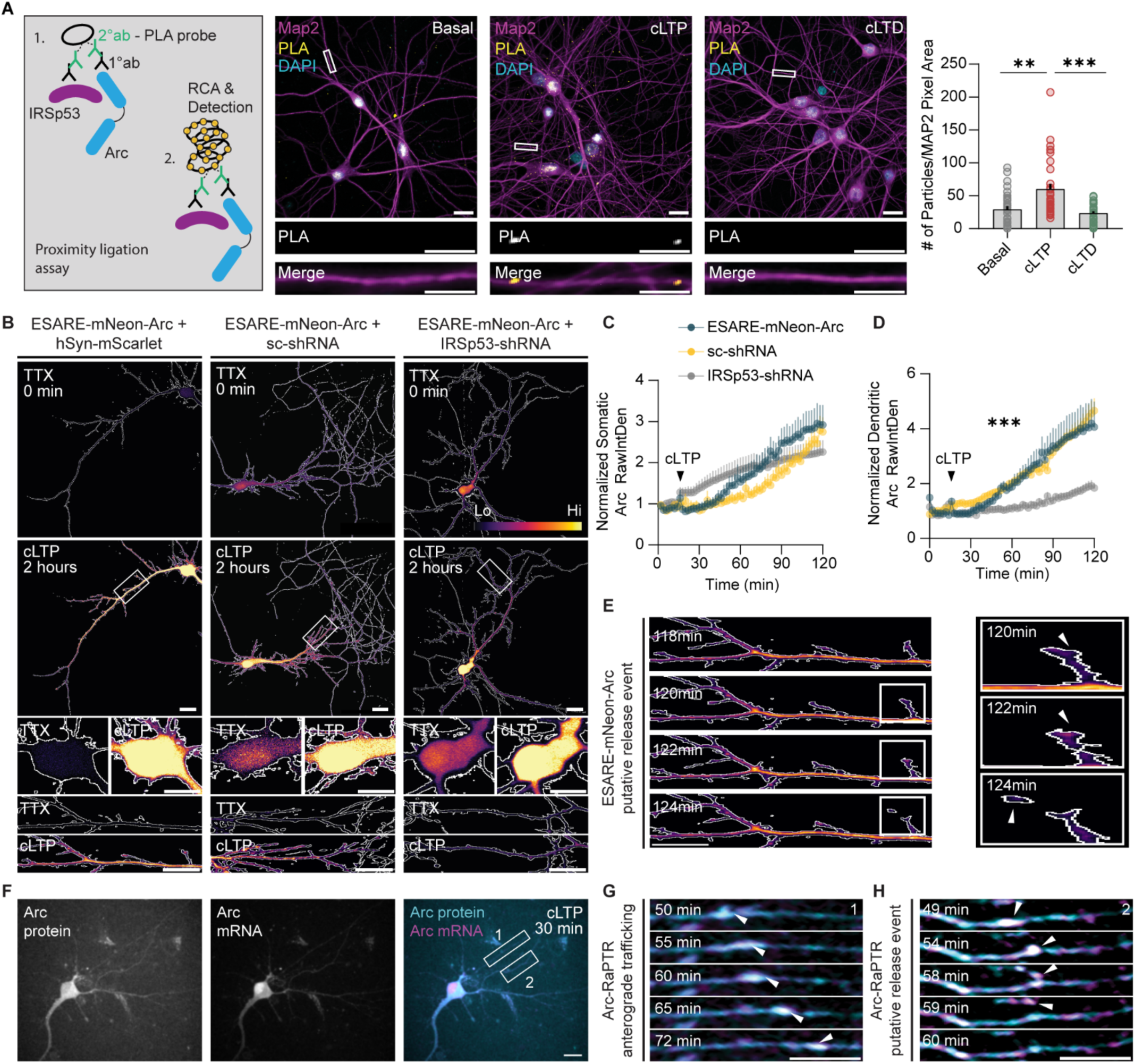
LTP induces IRSp53-dependent anterograde trafficking and release of Arc from dendrites. **A**. *cLTP induces Arc and IRSp53 interactions.* Proximity Ligation Assay. Representative images of primary cultured rat hippocampal neurons (DIV15) under basal, cLTP or cLTD conditions. cLTP, but not cLTD, increases Arc-IRSp53 puncta (basal: n=31, cLTP: n=28, cLTD: n=33 neurons) **B-D**. *cLTP induces IRSp53-dependent Arc anterograde trafficking into dendrites.* Representative max projection images of neurons in TTX prior to cLTP (0 mins) and 2 hrs post cLTP, co-transfected with ESARE-mNeon-Arc+hSyn-mScarlet or ESARE-mNeon-Arc treated with mScarlet-scrambled-shRNA (sc-shRNA) or mScarlet-IRSp53-shRNA. The cell outline was generated from an mScarlet mask and Arc expression is shown using the inferno LUT. **(C)**. Somatic Arc expression increases after cLTP induction in all conditions (black arrow=cLTP). **(D)**. Dendritic Arc levels increase after cLTP but not in IRSp53-shRNA treated neurons. Five 40µm dendritic segments were measured per neuron (n=5 neurons from 3 independent cultures). **E**. *Arc is released from filopodia-like protrusions.* Representative image of ESARE-mNeon-Arc in a dendrite, showing a putative release event after cLTP. **F-H**. *Arc protein and mRNA traffic together and are released from dendrites.* Representative image of Arc-RaPTR (RNA and Protein Translation Reporter), showing Arc protein and mRNA expression 30 min after cLTP induction. **(G)**. Time-lapse images of Arc protein and mRNA co-trafficking in dendrites after cLTP induction. **(H)**. Time-lapse images of Arc protein and mRNA in dendrites, showing a putative release event. Data are the mean ± s.e.m. One-way ANOVA with Dunn’s multiple comparison test (**A**) and Two-way ANOVA with Šídák’s multiple comparison test (**C-D**). \*\*\**P*<0.001, \*\*\*\**P*<0.0001. Scale bar=20µm. Dendritic scale bar=5µm.

**Supplementary Figure 2.**
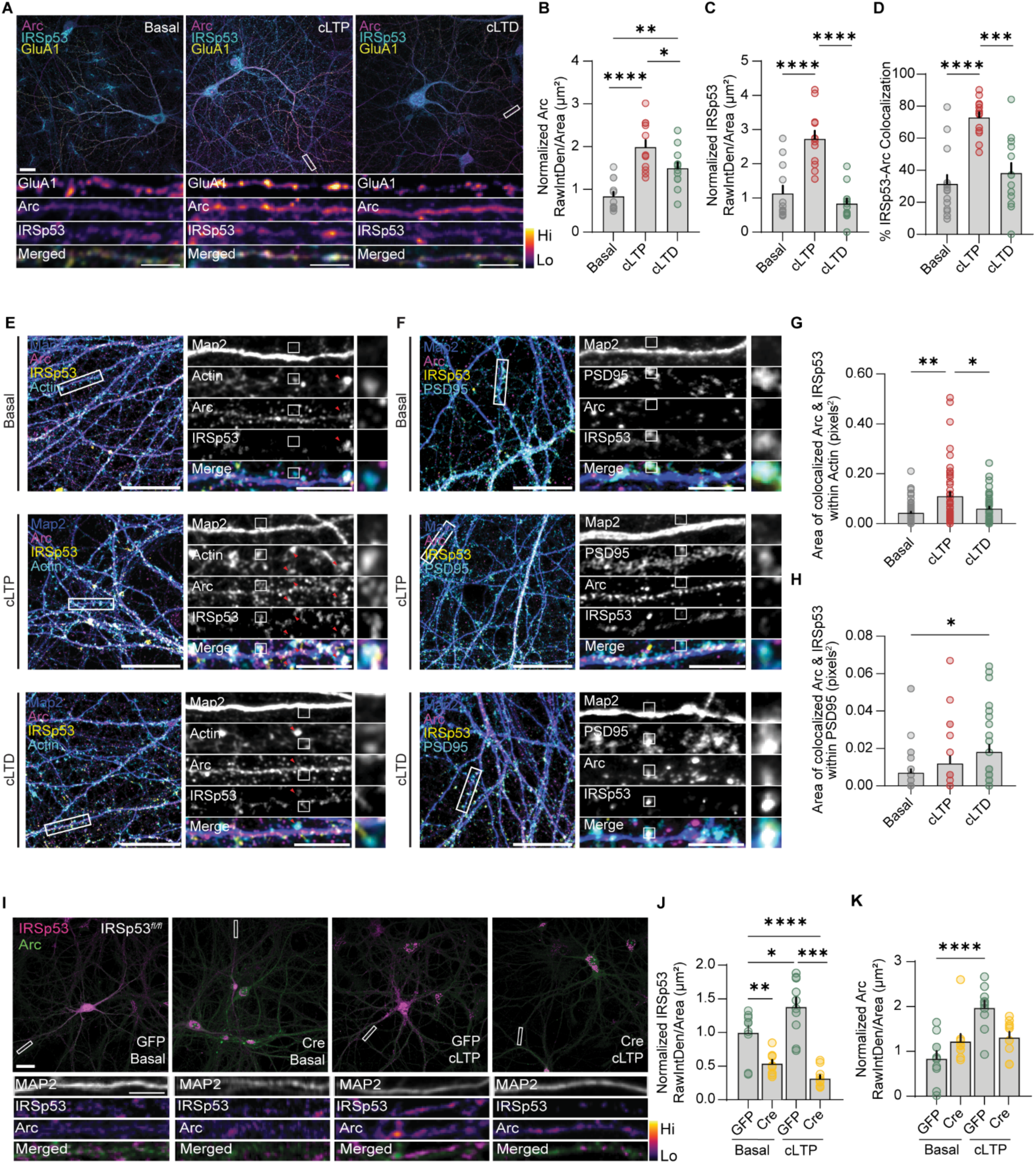
LTP induces Arc-IRSp53 colocalization in actin-rich extra-synaptic domains and release from dendrites. **A-D.** *cLTP induces Arc and IRSp53 colocalization in dendrites*. Representative images of primary cultured rat hippocampal neurons (DIV16) under basal, cLTP or cLTD conditions labeled with surface GluA1, and fixed/stained for Arc and IRSp53. cLTP and cLTD induce an increase in expression of Arc (**B**), but only cLTP induces an increase in IRSp53 expression (**C**) and an increase in Arc-IRSp53 colocalization (**D**). Arc and IRSp53 have increased colocalization after cLTP induction. Three 20µm dendritic segments were measured per neuron (n=14 neurons). **E-H**. *cLTP induces Arc and IRSp53 colocalization in actin-rich extra-synaptic areas in dendrites.* Representative images of primary hippocampal rat neurons imaged with a Zeiss Airyscan pseudo-super resolution module under basal, cLTP or cLTD conditions. 1 hour post cLTP/cLTD neurons were labeled with MAP2, Arc, IRSp53 and actin or PSD-95 (**F**). Arc and IRSp53 colocalize in actin-rich extra-synaptic regions following cLTP, but not in basal or cLTD conditions (**G**). After cLTD, Arc protein localizes with PSD-95 at synapses (**H**) (n=50 dendritic ROIs from 10 neurons). **I-K**. *IRSp53 is required for cLTP-induced Arc dendritic expression*. Representative images of *IRSp53^flx/flx^* primary cultured mouse hippocampal neurons under basal or cLTP conditions. Neurons were transduced with GFP or GFP-Cre lentivirus at DIV10 and neurons were fixed at DIV16. **(J).** Cre transduction decreases dendritic IRSp53 levels under basal conditions and prevents an increase in IRSp53 levels after cLTP induction. Three 20µm dendritic segments were measured per neuron (n=10 neurons). **(K)**. Cre-mediated knockdown of IRSp53 prevents an increase in dendritic Arc levels after cLTP. Three 20µm dendritic segments were measured per neuron (n=10 neurons). Data are the mean ± s.e.m. One-way ANOVA with Tukey’s multiple comparison test. \**P*<0.05, \*\**P*<0.005, \*\*\**P*<0.0005, \*\*\*\**P*<0.0001. Scale bar=20µm. Dendritic scale bar=5µm.

**Supplementary Figure 3.**
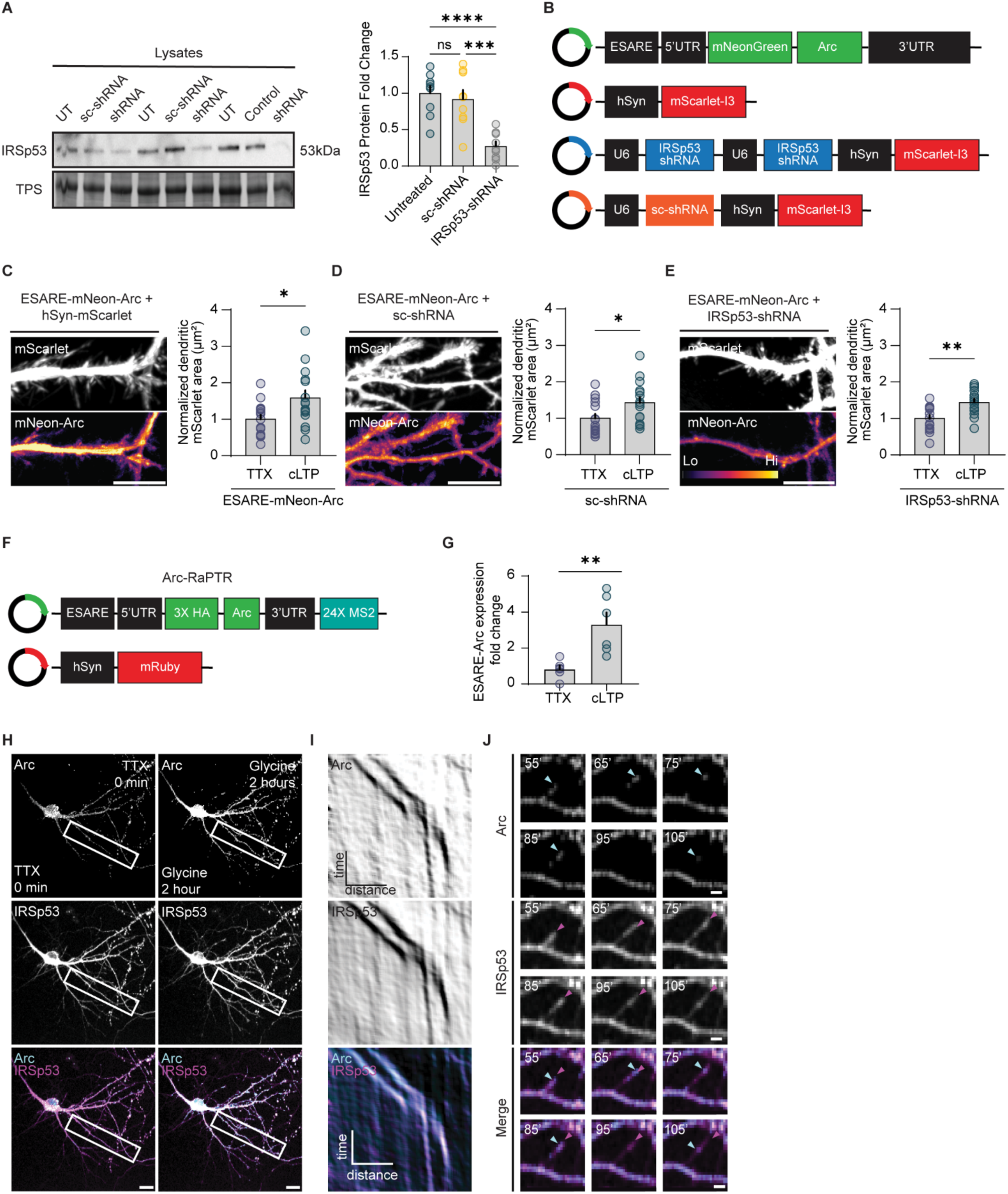
Validation of IRSp53 knockdown and effects on live cell dynamics. **A.** *Validation of IRSp53-shRNA knock-down.* Western blot and quantification of cell lysates from primary cortical neurons, showing a significant decrease in IRSp53 levels using IRSp53-shRNA. **B**. *Constructs used for live neuronal imaging experiments.* shRNAs targeting 2 separate exons were used for the IRSp53-shRNA construct. **C-E**. *IRSp53 knock-down does not affect cLTP-induced dendritic filopodia*. Representative time-lapse max projection images showing filopodial-like extensions following cLTP induction in (**C**) ESARE-mNeon-Arc, (**D**) ESARE-mNeon-Arc+IRSp53 sc-shRNA co-transfected, and (**E**) ESARE-mNeon-Arc+IRSp53 shRNA co-transfected dendrites. **F.** *Schematic of Arc-RaPTR (RNA and Protein Translation Reporter) and the mRuby cell filler plasmids.* **G**. *Arc-RaPTR expression increases after cLTP induction.* **H-I**. *Arc protein and mRNA traffic together in dendrites.* Representative max projection images of neurons co-transfected with the Arc-RaPTR reporter and a C-terminally tagged mRuby-IRSp53 immediately or 2 hours after cLTP induction. **(I)** Kymograph showing anterograde co-trafficking of Arc and IRSp53 in dendrites following cLTP. Time is displayed on the y-axis (bar=25 min.) and distance on the x-axis (bar=30µm). **J.** *cLTP induced Arc and IRSp53 dendritic filopodia*. Dendritic ROI showing an IRSp53 labeled filopodial-like extension (magenta) with Arc protein at the leading tip (cyan arrow). There is rapid loss of Arc signal from this extension. Data are the mean ± s.e.m. One-way ANOVA with Tukey’s multiple comparison test (**A**) and unpaired t-test with Welch’s correction (**C-E**, **G)**. \**P*<0.05, \*\**P*<0.005, \*\*\**P*<0.0005, \*\*\*\**P*<0.0001, ns=not significant. Scale bar=20µm. Dendritic scale bar=5µm.

Given these data, we investigated the dynamics of Arc-IRSp53 trafficking in living neurons. Primary rat hippocampal neurons were co-transfected with ESARE (the minimal Arc promoter and upstream activity-dependent transcription factor binding sites that mimic endogenous Arc expression^34^) Arc tagged with mNeonGreen (mNeon) and either hSyn-mScarlet, an IRSp53-shRNA construct to knock-down endogenous IRSp53 (Figure S3A), or a scrambled shRNA (Figure S3A-B). cLTP increased somatic Arc expression in all conditions (Figure 2B-C). However, dendritic Arc expression after cLTP was significantly lower in IRSp53-shRNA co-transfected cells (Figure 2B, D). Similarly, cLTP did not increase dendritic Arc expression in *IRSp53^flx/flx^* neurons from an IRSp53 floxed mouse line^35^ transduced with Cre (Figure S2I-K). cLTP induced anterograde trafficking of Arc protein from the cell body into dendrites (Figure 2B-C and Supplementary Movie 1), which was abrogated in IRSp53-shRNA transfected neurons (Figure 2B and Supplementary Movie 2). IRSp53 is known to regulate dendritic filopodia and spines^18,19^. After cLTP induction, transient filopodial-like protrusions were observed in all conditions, with no significant differences in their overall number (Figure S3C-E). In control neurons, many of these protrusions contained Arc puncta that rapidly disappeared (Figure 2E and Supplementary Movie 3; the white arrow shows the filopodia location, and the orange arrow shows the putative release event), which may be release events. In contrast, neurons expressing IRSp53-shRNA still generated filopodia but showed markedly reduced Arc trafficking into dendrites and lacked Arc-filled filopodia (Figure S2B, D and S3E). These data suggest that IRSp53 facilitates Arc trafficking into dendrites. To confirm this, we imaged Arc-RaPTR (RNA and Protein Translation Reporter), which allows imaging of both Arc protein and mRNA (Figure S3F). Hippocampal neurons were transfected with Arc-RaPTR alone or co-transfected with IRSp53-mRuby (Figure S3F). cLTP induced an increase in Arc expression and the trafficking of Arc-IRSp53 puncta in dendrites (Figure S3G-I and Supplementary Movie 4). cLTP induced transient formation of filopodial-like protrusions in dendrites containing Arc and IRSp53, followed by a rapid loss of Arc signal in these protrusions (Figure S3J and Supplementary Movie 4). We also observed co-trafficking of both Arc protein and mRNA in dendrites (Figure S3F-G and Supplementary Movie 5), as well as putative release events (Figure S3H and Supplementary Movie 6). These observations show that LTP induces the co-trafficking of Arc protein, *Arc* mRNA, and IRSp53 in dendrites, where Arc release occurs from filopodia-like protrusions.

### IRSp53 mediates Arc extracellular vesicle release

Retroviruses are released from cells in membrane-enveloped particles and IRSp53 is critical for HIV release^23,24^. To determine whether IRSp53 regulates Arc release, we used a split-nano luciferase system^36^, where we cloned the small HiBiT tag (11 amino acids) onto the Arc C-terminus (Figure 3A), which does not interfere with capsid formation (Figure S4A). We transfected HEK293T cells with Arc-HiBiT alone, Arc-HiBiT+IRSp53, or Arc-HiBiT+GFP. Cell lysate and media samples were collected to measure luciferase luminescence, which occurs after HiBiT complementation with Large-bit (LgBiT) (Figure 3A). When IRSp53 is co-expressed with Arc, release is enhanced, whereas GFP co-expression has no effect (Figure 3B). To determine whether Arc-HiBiT released into the media is enclosed within EVs, we measured luminescence in the presence or absence of a membrane-emulsifying detergent. Luminescence was only observed in detergent-treated media, suggesting that Arc-HiBiT is protected within the EV lumen (Figure 3C). Using SEC and nanoparticle tracking, we find that IRSp53 facilitated Arc protein release in particles with the size and density of EVs (Figure 3D-E). IRSp53 did not facilitate Arc^Δ20–40^ release (Figure 3F), consistent with Arc^Δ20–40^ being unable to bind IRSp53 (Figure 1C). Additionally, we used U87 cells transduced with IRSp53 under the control of a doxycycline-inducible promoter. Cells were treated with increasing concentrations of doxycycline, which increased IRSp53 expression in cell lysates and increased Arc release into the media (Figure S4B). IRSp53 may facilitate Arc release through its SH3 domain via downstream interactions with actin effector proteins, such as WAVE2, N-WASP, and Dynamin1^37^. An IRSp53 SH3 point mutant that blocks these interactions, IRSp53^I403P^, does not enhance Arc release in HEK293T cells (Figure 3F) but is still capable of binding to Arc protein and facilitating Arc capsid assembly (Figure S4C-D). Furthermore, Endophilin, an N-BAR domain-containing protein known to interact with Arc^4^ did not facilitate Arc release from HEK293T cells (Figure S4E). IRSp53 overexpression also enhances Arc but not Arc^Δ20–40^ release in neurons, whereas IRSp53^I403P^ has no effect (Figure 3G).

**Figure 3.**
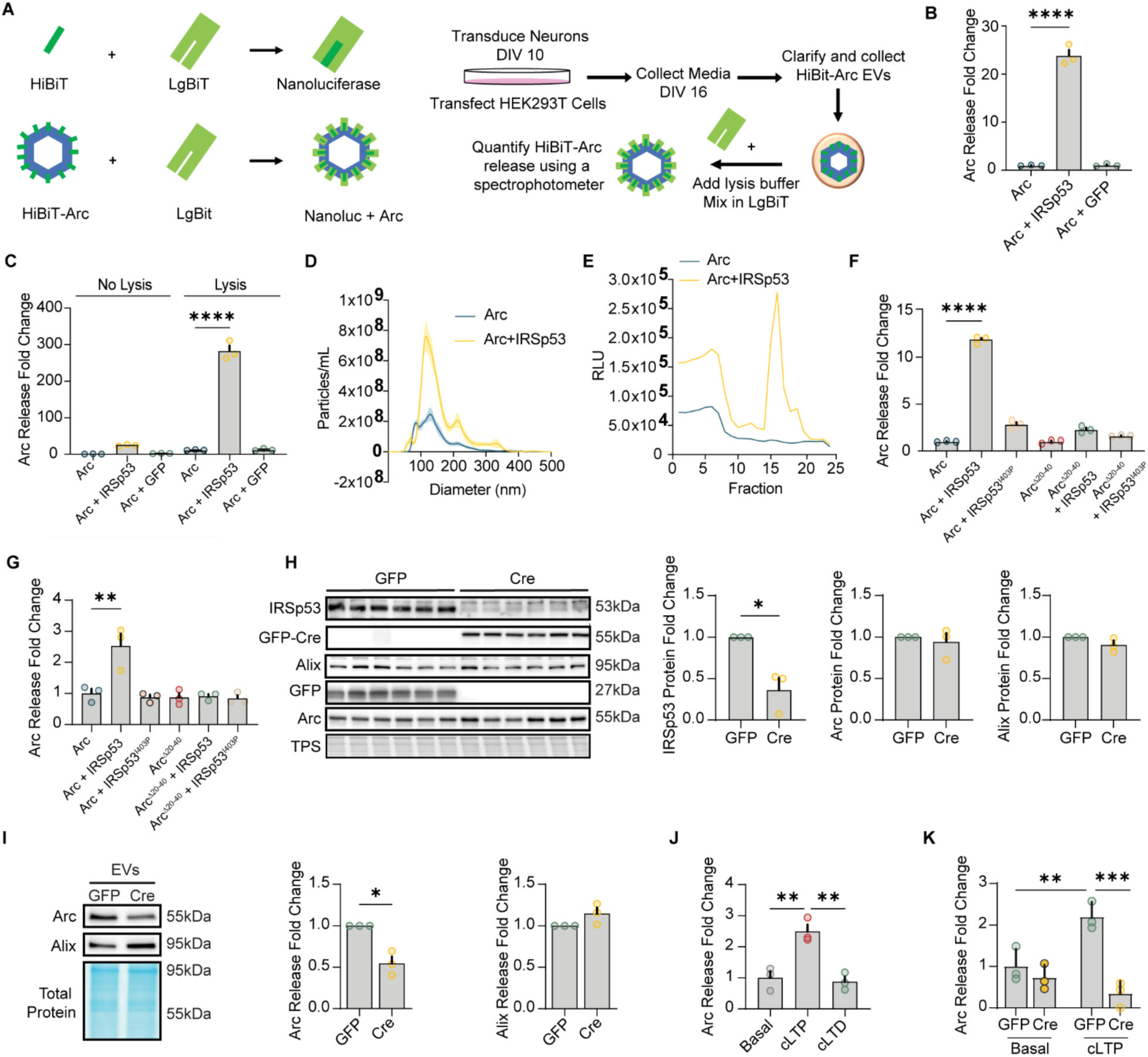
IRSp53 mediates Arc extracellular vesicle release. **A**. *HiBiT split nanoluciferase assay.* Neurons were transduced or HEK293T cells were transfected and EVs were collected, lysed, and purified and LgBiT added for luminescence quantification. **B**. *IRSp53 facilitates Arc release in HEK293 cells*. HEK293T cells were transfected with Arc, Arc+IRSp53, or Arc+GFP. Percentage Arc release was calculated by dividing the total luminescence in the media by the sum of total luminescence in the cell lysate and total luminescence in the media (n=3 cultures). **C**. *Arc is released from HEK293 cells in EVs.* Luminescence was undetectable in the absence of lysis, indicating that Arc-HiBiT is protected within vesicles. **D**. *Arc is released in small EVs.* The size of Arc EVs was determined using nanoparticle tracking analysis (NTA). Particle tracks (n=3 independent EV samples) are depicted as averages (blue and yellow line traces) with ± SEM depicted as shaded areas. **E**. *IRSp53 facilites Arc EV release.* Equilibrium density gradient isolation was used to isolate Arc EVs from HEK293T cell culture media. The density in fractions 15-17 was between 1.10 and 1.15 grams/mL, consistent with the known densities of EVs. **F**. *IRSp53 enhanced Arc release is specific and requires its SH3 domain.* IRSp53 does not enhance the release of Arc^Δ20–40^ in HEK293T cells (n=3 cultures). **G**. *IRSp53-dependent enhancement of Arc release requires the interaction with actin machinery (IRSp53^I403P^) and direct protein interaction with Arc (Arc^Δ20–40^) in neurons* (n=3 cultures). **H**. *Cre induces IRSp53 knock-down in cells*. Representative western blot of cell lysate protein expression from primary cortical neurons transduced with GFP or GFP-Cre. Cre expression decreases IRSp53 levels while Arc and Alix are unaffected. **I**. *IRSp53 is required for Arc EV release*. Representative western blot of Alix or Arc in EVs released from primary cortical neurons. Arc and EV expression was normalized to GFP transduced neurons. Arc release is significantly reduced (n=3 independent cultures) **J**. *cLTP increases Arc-HiBiT release.* **K**. *cLTP-induced Arc release is IRSp53-dependent.* Cre-mediated knockdown of IRSp53 prevents LTP-driven Arc release. Each data point represents one well from a 6-well plate (technical replicates) and are representative of 3 independent experiments. Data are the mean ± s.e.m. One-way ANOVA with Dunnett’s multiple comparison test (**B,C,F,G,J**), unpaired t-test with Welch’s correction (**H,I**), and Two-way ANOVA with Fisher’s LSD (**K**). \**P*<0.05, \*\**P*<0.005, \*\*\**P*<0.0005, \*\*\*\**P*<0.0001, ns=not significant.

**Supplementary Figure 4.**
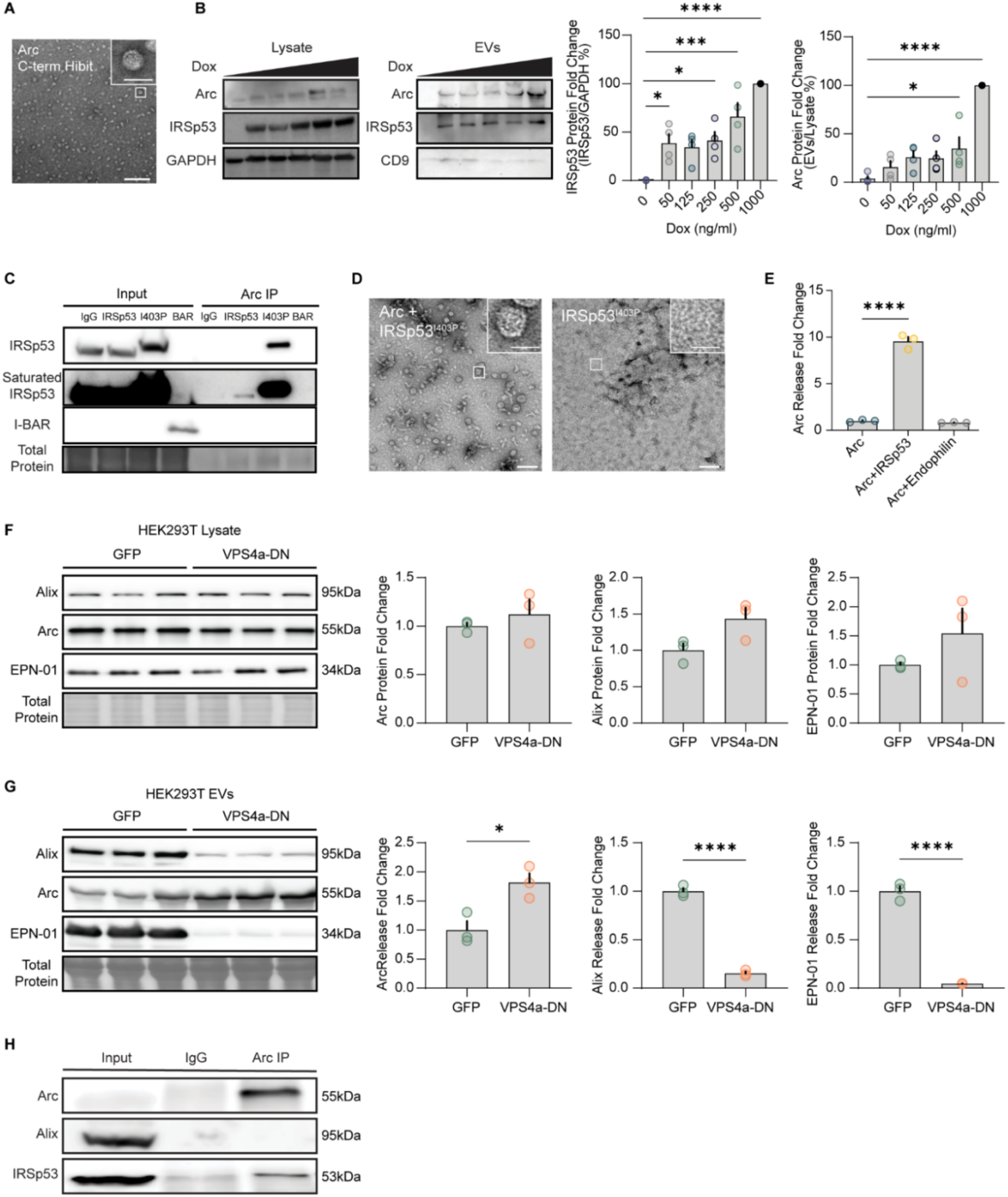
IRSp53-dependent Arc capsid assembly and release mechanisms. **A**. *Arc tagged with HiBiT at the C-terminus does not interfere with capsid formation.* Scale bar=200 nm. **B**. *IRSp53 increases Arc EV release.* Western blot of U87 cells transduced with a doxycycline-inducible IRSp53 virus. Increasing doxycycline increases IRSp53 expression and Arc release in EVs (n=4 independent cultures). **C.** *Arc-IRSp53 binding interactions*. HEK293T cells were transfected with Arc+IRSp53 or IRSp53^I403P^ or the IRSp53 I-BAR domain. Arc binds to both IRSp53 and IRSp53^I403P^, but not the IRSp53 I-BAR domain. **D**. *IRSp53^I403P^ facilitates Arc capsid assembly similar to WT IRSp53.* **E**. *Endophilin, an N-BAR containing protein that also interacts with Arc, does not affect Arc release in HEK293T cells* (n=3 independent cultures). **F-G**. *Arc release does not require ESCRT activity.* Representative western blot of lysate from HEK293T cells transfected with GFP or a VPS4a dominant-negative (VPS4a-DN) construct. The expression of VPS4a-DN does not significantly affect the expression of Arc, Alix, or a EPN-01 protein nanocage^40^ in the cell lysate. **(G)**. Representative western blot of EVs collected from the media. VPS4a-DN expression increases Arc release while reducing both Alix and EPN-01 release. **H**. Arc *does not bind Alix in mouse brain.* A one month old WT mouse was enriched for six hours, and the forebrain was collected. Arc was IPed and the sample was blotted for Alix and IRSp53. Data are the mean ± s.e.m. One-way ANOVA with Dunnett’s multiple comparison test (**B, E**), unpaired t-test with Welch’s correction (**F, G**). \**P*<0.05, \*\*\**P*<0.0005, \*\*\*\**P*<0.0001, ns=not significant.

We next determined whether Arc release is dependent on the ESCRT pathway, which is known to be important for exosome and retrovirus secretion^38,39^. We transfected HEK293T cells with a VPS4a dominant-negative (VPS4a-DN) construct that blocks ESCRT-dependent processes^40^. VPS4a-DN expression did not affect cell lysate expression of Arc, Alix (a canonical EV marker and ESCRT protein), or the protein nanocage EPN-01 that contains an HIV Gag P6 ESCRT binding motif^40^ (Figure S4F). VPS4a-DN expression significantly reduced the release of both Alix and EPN-01, but Arc expression was paradoxically increased (Figure S4G). Moreover, Arc and Alix do not co-IP from mouse cortex (Figure S4H). This shows that Arc release does not occur through conventional exosome or retrovirus budding mechanisms, consistent with a lack of a conserved ESCRT P6 binding motif.

To determine whether IRSp53 plays a role in the release of endogenous Arc in EVs from neurons, we used the IRSp53 floxed mouse line^35^ and generated *IRSp53^flx/flx^* primary cultured neurons. Lentiviral expression of GFP-Cre for six days (DIV10-16) resulted in a significant reduction in IRSp53 expression in cell lysates as compared with GFP expression, but no change in Arc or Alix expression (Figure 3H). In Cre-expressing neurons, we observed a significant decrease in Arc EV release, but no effect on Alix levels in EVs (Figure 3I). cLTP increased Arc release, as compared to basal release and cLTD treatment (Figure 3J). However, cLTP had no effect on Arc release in *IRSp53^flx/flx^* neurons that were transduced with Cre (Figure 3K). These results show that IRSp53 is critical for activity-dependent release of Arc EVs from neurons.

### Arc EVs induce the loss of surface AMPARs and weaken synapses in recipient neurons

What is the functional significance of Arc release in EVs? Arc has a well-established role in trafficking AMPA-type glutamate receptors (AMPARs)^4,41^. Thus, we hypothesized that Arc EVs may alter synapse function (Figure 4A). To complement our LTP studies in WT rat neurons, we generated an Arc knock out (KO) rat using CRISPR (Figure S5A-B). We isolated EVs from WT or Arc KO rat cortical neurons and added 50µg of total EVs to Arc KO rat hippocampal neurons for 4 hours, followed by a live antibody feeding assay to label the surface AMPAR subunit GluA1^8^. Neurons treated with WT EVs had a significant reduction in surface GluA1 levels, while neurons treated with Arc KO EVs had no difference in surface GluA1 levels (Figure 4B). The loss of surface AMPARs may result from the delivery of Arc protein in EVs or *Arc* mRNA in capsids, which could then be locally translated in the recipient neuron. To test this, we transduced Arc KO recipient neurons with an Arc-specific shRNA lentivirus, which blocks the translation of *Arc* mRNA (Figures 4A and S5C). Treatment with Arc-shRNA did not affect surface AMPAR levels (Figure 4C). Neurons transduced with Arc-shRNA or sc-shRNA showed no changes in surface GluA1 when incubated with WT EVs, while neurons transduced with scrambled shRNA showed decreased surface GluA1 levels after WT EV incubation (Figure 4C). To determine whether Arc EVs directly affect synaptic function, we measured mini excitatory postsynaptic currents (mEPSCs), using whole-cell voltage clamping, in cultured Arc KO rat hippocampal neurons that were incubated with WT or Arc KO EVs. WT EVs significantly reduced mEPSC amplitude, but not frequency (Figure 4D-E), as compared to Arc KO EVs. These data show that Arc EVs reduce synaptic strength via delivery of *Arc* mRNA and subsequent local translation that induces a loss of surface AMPARs.

**Figure 4.**
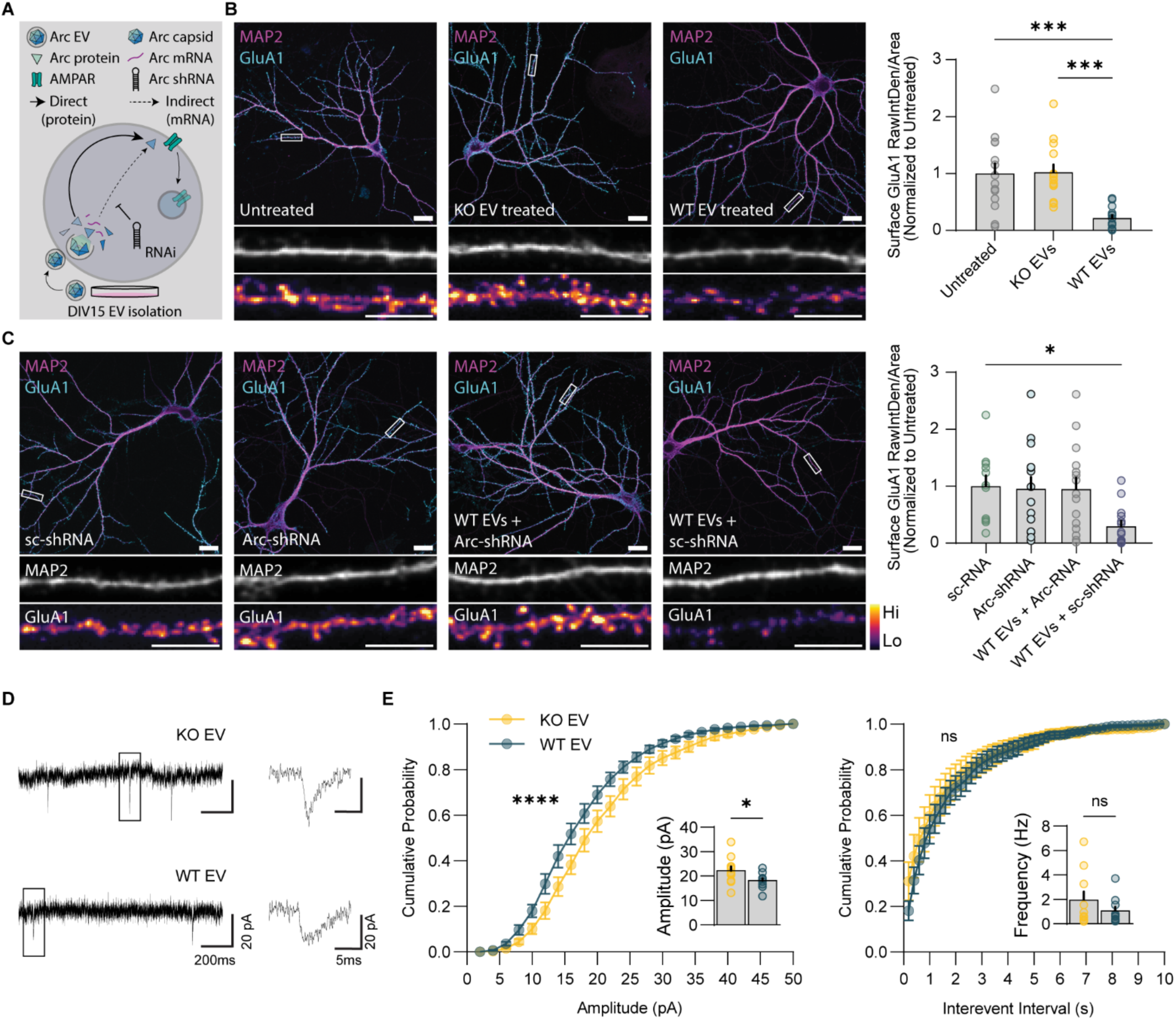
Arc EVs weaken synaptic strength by decreasing surface AMPA receptors. **A**. *Experimental paradigm to test whether Arc protein or mRNA delivery via EVs is important for functional effects on AMPARs*. **B**. *Arc EVs decrease surface dencdritic GluA1 levels.* Representative images of DIV 14 Arc KO primary rat hippocampal neurons incubated with WT or KO EVs for 4 hours and labelled for surface AMPARs. WT EVs result in a significant loss of surface AMPARs, while KO EVs had no effect. Three 20µm dendritic segments were measured per neuron (n=15 neurons). **C**. *Arc EV-induced loss of surface GluA1 requires Arc translation.* Representative images of neurons that were transduced with either scrambled shRNAs (sc-shRNA) or Arc-shRNAs at DIV 10 and treated with WT EVs on DIV 15. Neurons transduced with Arc-shRNA or sc-shRNA showed no loss of surface AMPAR with WT EV incubation, as compared with a significant loss observed in neurons transduced with sc-shRNA. Three 20µm dendritic segments were measured per neuron (n=15 neurons). **D-E**. *Arc EVs decrease synapse strength.* Representative mEPSC traces of neurons treated with either KO or WT EVs. **(E).** Cumulative probability plots of mEPSC amplitude and frequency after treatment with KO or WT EVs, showing a significant decrease in mEPSC amplitude but no effect on frequency. Data are the mean ± s.e.m. One-way ANOVA with Dunnett’s multiple comparison test (**B, C**), two-way ANOVA with Šídák’s multiple comparisons test and unpaired t-test with Welch’s correction (**E**). \**P*<0.05, \*\*\**P*<0.0005, \*\*\*\**P*<0.0001, ns=not significant. Scale bar=20µm. Dendritic scale=5µm.

**Supplementary Figure 5.**
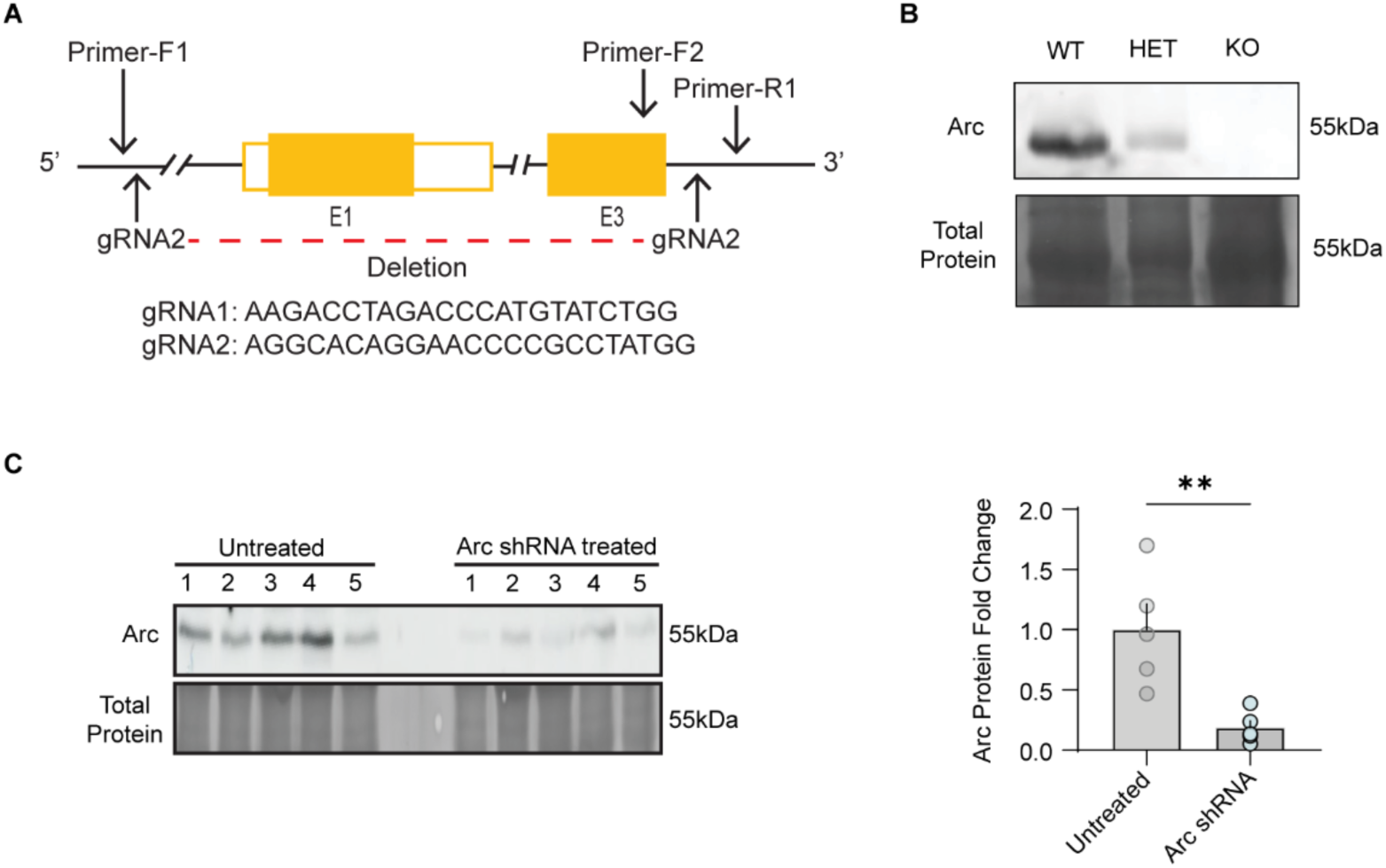
Validation of Arc KO rat and Arc shRNA. **A**. *Arc rat KO design.* The CRISPR design strategy for excising the entire *Arc* gene in Long-Evans rats. Embryos were injected with gRNAs and dCAS9 before being implanted in a dam. **B**. *Arc rat KO validation.* Western blot of rat forebrain tissue collected from WT, heterozygous, and Arc KO rats showing a complete loss of Arc protein in the KO. **C**. *Arc-shRNA validation.* Western blot of primary cultured neuronal lysates obtained from untreated cells, or cells transduced with Arc shRNA lentivirus. Arc shRNA expression significantly reduced Arc protein levels in WT neurons (n=5 10cm dishes from one culture). \*\**P*<0.005. Unpaired t-test with Welch’s correction.

### LTP induces intercellular Arc transfer

We reasoned that Arc EV uptake may occur in neighboring neurons. We sparsely expressed either GFP alone or GFP+ESARE-Arc in rat Arc KO primary hippocampal neurons. ESARE-Arc expression increases after cLTP, to similar levels of endogenous Arc (Figure S6A-C), while PRK5-Arc plasmids that contain the CMV promoter do not respond to neuronal activity (Figure S6A-E). Both cLTP and cLTD increased ESARE-Arc expression in transfected cells compared to basal conditions (Figure 5A-C). However, only cLTP increased Arc levels in neighboring non-transfected dendrites (Figure 5A, D). Significant cell-to-cell transfer of Arc was not observed in recipient neuron cell bodies (Figure 5A, E), although this may be due to higher non-specific background staining in cell bodies than dendrites. Constitutive Arc expression in mouse primary hippocampal neurons also resulted in a significant increase in dendritic Arc staining in neurons neighboring Arc-transfected neurons, as compared to neighbors of GFP-transfected neurons (Figure S6F-H). Together, these data show that LTP and high Arc expression result in the transfer of Arc protein into nearby dendrites.

**Figure 5.**
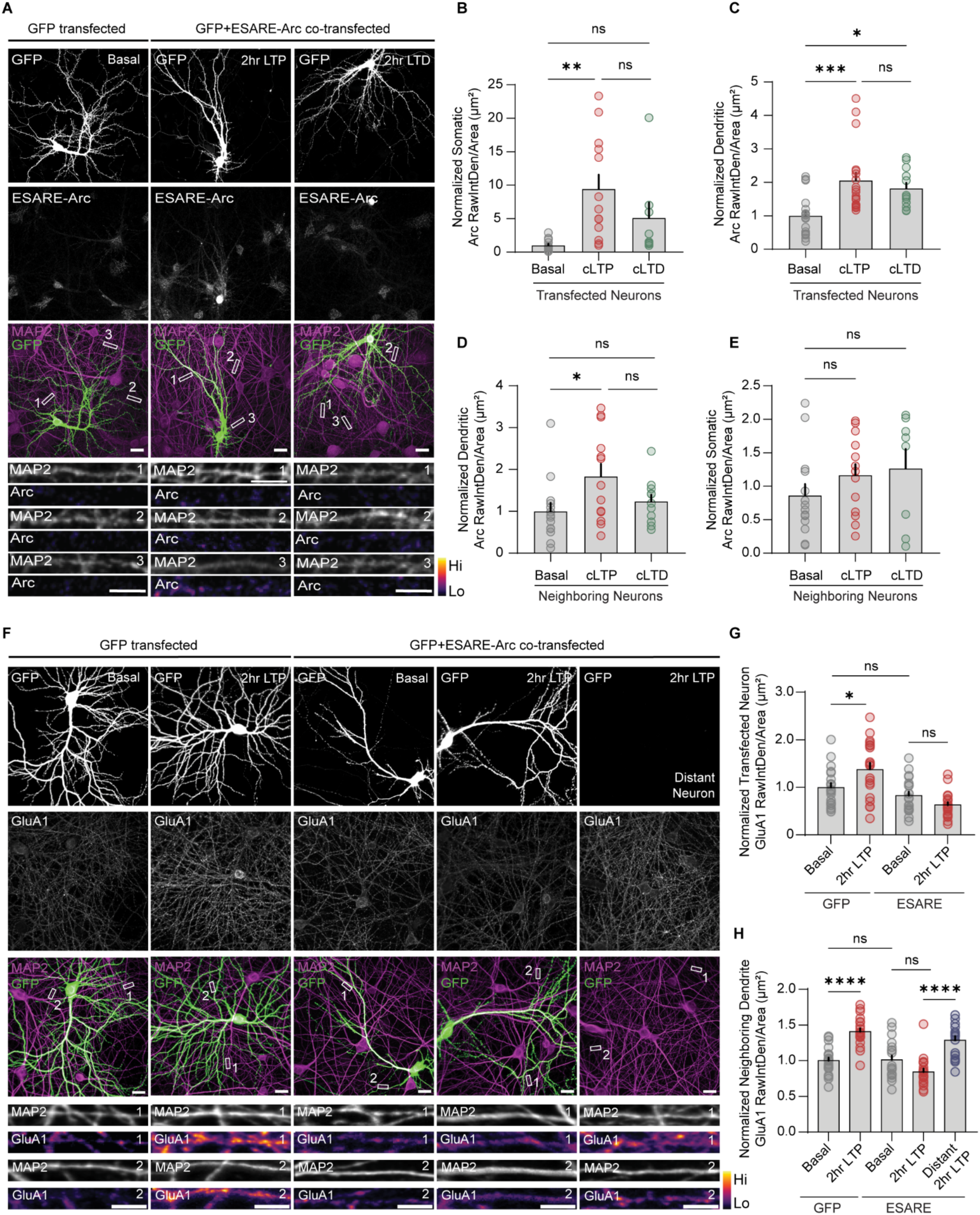
LTP induces Arc intercellular transfer. **A-E**. *Arc transfer to neighboring untransfected dendrites was observed after cLTP but not cLTD*. Representative images of GFP and GFP+ESARE-Arc co-transfected primary cultured hippocampal Arc KO rat neurons in basal, cLTP, or cLTD conditions. Neurons were labeled for Arc, GFP, and MAP2 at DIV15. Quantification of raw integrated density/area normalized to basal conditions for **(B)** somatic Arc levels in transfected neurons, (**C**) dendritic Arc levels in transfected neurons, (**D**) somatic Arc levels in neighboring neurons, and (**E**) dendritic Arc levels in neighboring dendrites (basal, cLTP, and cLTD dendrites, n=8-12 neurons, three dendritic segments/neuron). **F-H**. *Arc transfer prevents LTP-induced increases in surface GluA1 levels.* (**F**) Representative images of GFP and GFP+Esare-Arc co-transfected primary cultured Arc KO rat neurons in basal or cLTP conditions. (**G**) ESARE-Arc expression had no effect on AMPAR levels in basal conditions. cLTP induction increases surface AMPAR levels in GFP-transfected cells, whereas no difference was observed in ESARE-Arc transfected neurons (n=20 neurons, five dendritic segments/neuron). (**H**) cLTP resulted in an increase in AMPAR levels in dendrites neighboring GFP-transfected neurons. In contrast, cLTP had no effect on surface AMPAR levels in dendrites neighboring ESARE-Arc co-transfected neurons. cLTP induced an increase in surface AMPARs in dendrites greater than 400µm away from Arc transfected neurons. Five dendritic segments were measured per neuron (n=20 neurons per group). Data are the mean±s.e.m. One-way ANOVA with Tukey’s multiple comparison test (**B-E, G, H**).\**P*<0.05, \**P*<0.005, \*\*\**P*<0.0005, \*\*\*\**P*<0.0001, ns=not significant. Scale bare=20µm. Dendritic scale=5µm.

**Supplementary Figure 6.**
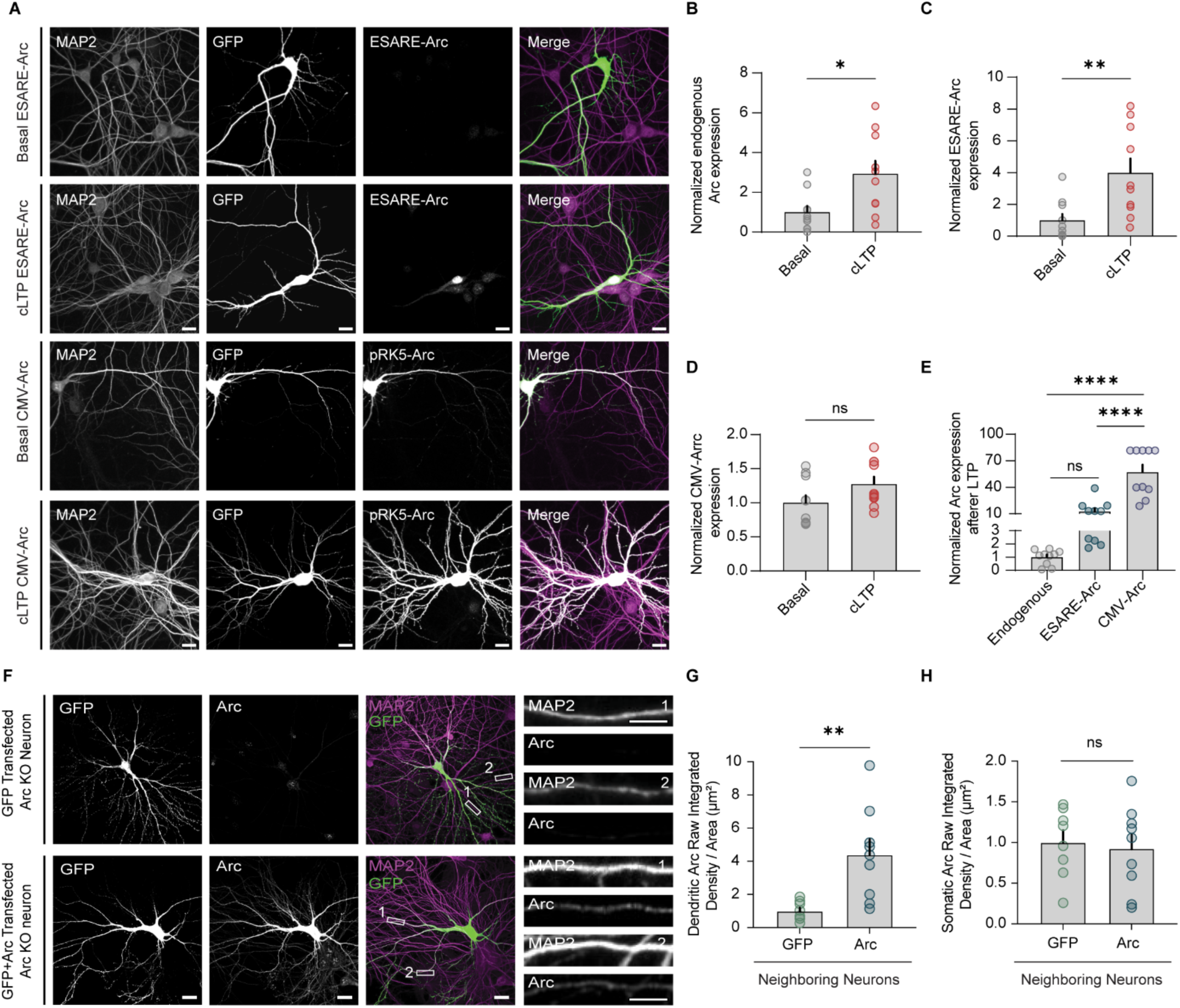
High constitutive expression of Arc results in intercellular Arc transfer to neighboring dendrites. **A-D**. *High Arc levels induce Arc transfer.* Representative images of primary WT rat hippocampal neurons co-transfected with GFP and ESARE-Arc or CMV-Arc in basal or cLTP conditions. cLTP results in an increase in expression for both endogenous Arc and ESARE-Arc. CMV-Arc levels do not change after cLTP and are considerably higher than endogenous and ESARE-Arc (n=10 neurons). Scale bar=20µm. cLTP induces endogenous (**B**) and ESARE-Arc (**C**) expression but not CMV-Arc (**D**). **E**. *CMV-Arc expression is significantly higher than endogenous and ESARE-Arc*. Image acquisition settings to visualize endogenous and ESARE-Arc saturate CMV-Arc pixels. **F.** Representative images of primary cultured Arc KO hippocampal neurons transfected with GFP and GFP+CMV-Arc, stained for Arc and MAP2 at DIV15. **G-H.** Arc levels in dendrites neighboring Arc-transfected neurons are significantly increased above GFP transfected/background antibody staining (n=8-12 neurons). Data are the mean ± s.e.m. One-way ANOVA with Tukey’s multiple comparison test (**E**) and unpaired t-test with Welch’s correction (**B-D, G, H**). \**P*<0.05, \*\**P*<0.01, \*\*\*\**P*<0.0001. Scale bar=20µm. Dendritic scale=5µm.

### Arc intercellular transfer regulates surface AMPA receptors

Given the effect of WT EVs on AMPAR levels and mEPSCs, we reasoned that intercellular Arc transfer could regulate AMPAR expression in neighboring neurons. To test this, we sparsely expressed ESARE-Arc in primary rat hippocampal Arc KO neurons and measured surface GluA1 levels in neighboring dendrites and cell bodies after cLTP. Under basal conditions, surface GluA1 levels were comparable between GFP-only and GFP+ESARE-Arc co-transfected neurons (Figure 5F-G), corroborating no significant over-expression artifacts. As expected, cLTP increased surface GluA1 in both GFP-transfected neurons (Figure 5F-G) and the untransfected neighboring neurons in the same coverslip (Figure 5H). In contrast, ESARE-Arc–expressing neurons showed no cLTP-induced increase in surface GluA1 (Figure 5F, G), and neighboring dendrites adjacent to these neurons also failed to show an increase in surface GluA1 (Figure 5A, H). Neurons located farther from ESARE-Arc–expressing cells in the same cover slip, however, had normal cLTP-induced increases in surface GluA1 (Figure 5A, H), indicating that the effect is spatially restricted. No significant changes were detected in neighboring cell bodies under any condition (Figure S7A).

Given that cLTP activates signaling cascades in all neurons and alters AMPAR levels in most neurons, we next asked whether Arc transfer alone could modulate AMPARs in the absence of LTP. We expressed CMV-Arc under basal conditions which resulted in a significant decrease in surface GluA1 in both Arc-transfected neurons and neighboring dendrites spatially proximal to Arc-transfected neurons (Figure S7B, D-E). No changes were observed in distant neurons on the same coverslip (Figure S7C, E). In contrast, GFP alone expressed in neurons had no effect on surface GluA1 in transfected neurons or neighboring dendrites (Figure S7B-E). Together, these results show that intercellular Arc transfer locally prevents cLTP induced increases in surface AMPARs in recipient neighboring dendrites, due to increased endocytosis and loss of surface GluA1, as observed with constitutive expression of Arc.

**Supplementary Figure 7.**
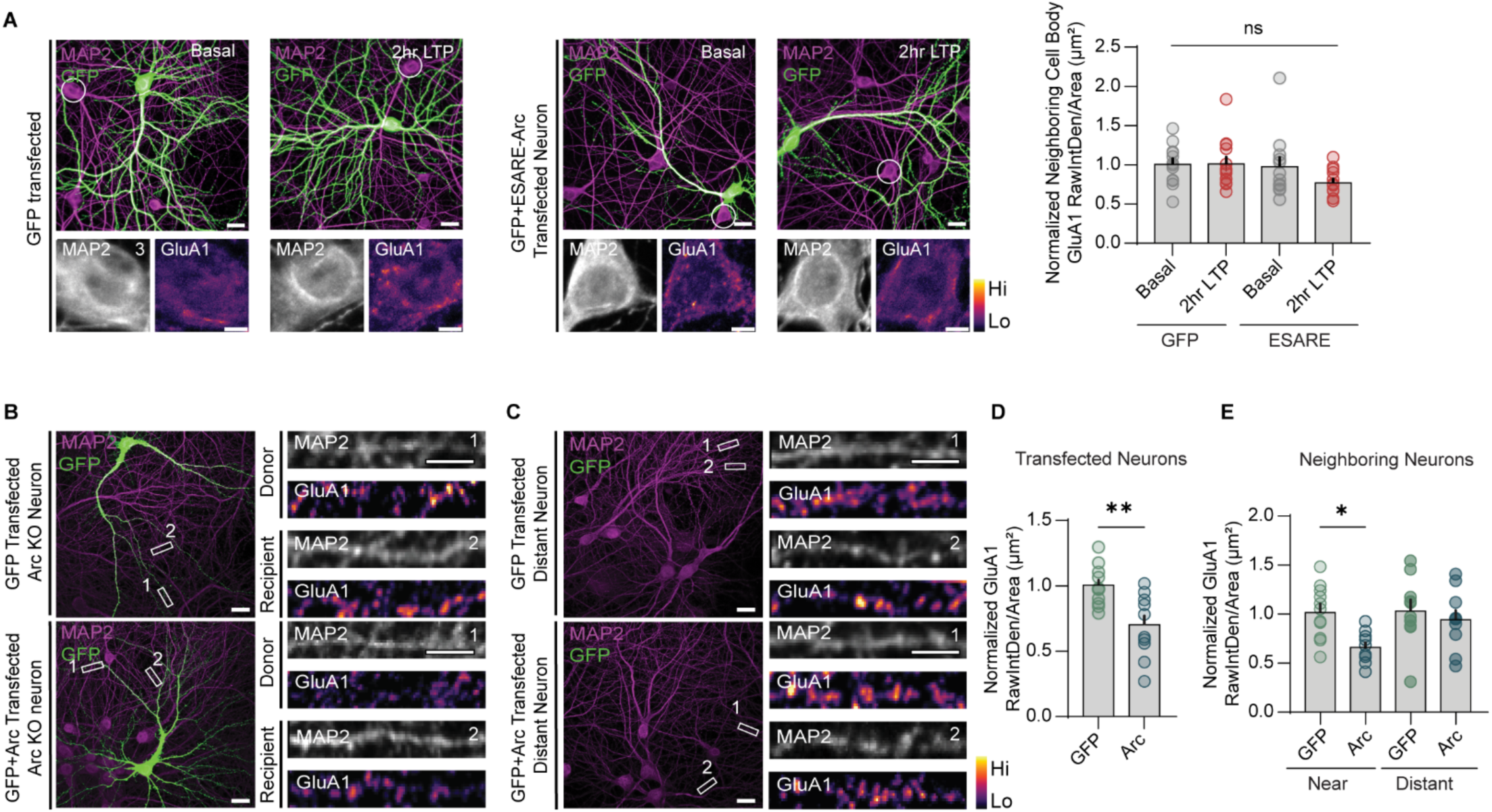
High constitutive expression of Arc results in the loss of surface AMPARs in neighboring dendrites. **A.** *cLTP does not induce significant differences in surface GluA1 in cell bodies.* Representative images of GFP or GFP+ESARE-Arc transfected Arc KO rat primary neurons in basal or cLTP conditions. No changes in somatic surface GluA1 levels were observed after cLTP induction (n=20 neurons). **B**. Representative images of a GFP and GCP+CMV-Arc co-transfected rat Arc KO primary hippocampal neurons stained for MAP2 and surface GluA1 at DIV15. **C**. Same as (**B**) except images were taken > 400µm from a transfected neuron in the same coverslip. **B-E.** *Arc decreases surface AMPAR expression in transfected cells* (**D**) *as well as neighboring non-transfected dendrites* (**E**). *No changes in GluA1 levels were observed in GFP neighboring dendrites or neurons further than 400µm* (**E**). Three dendritic segments were measured per neuron (n=10-12 neurons from 3 coverslips). Data are the mean ± s.e.m. One-way ANOVA with Tukey’s multiple comparison test (**A**) and unpaired t-test with Welch’s correction (**D, E**). \*\**P*<0.005, ns=not significant. Scale bar=20µm. Dendritic scale=5µm.

**Supplementary Figure 8.**
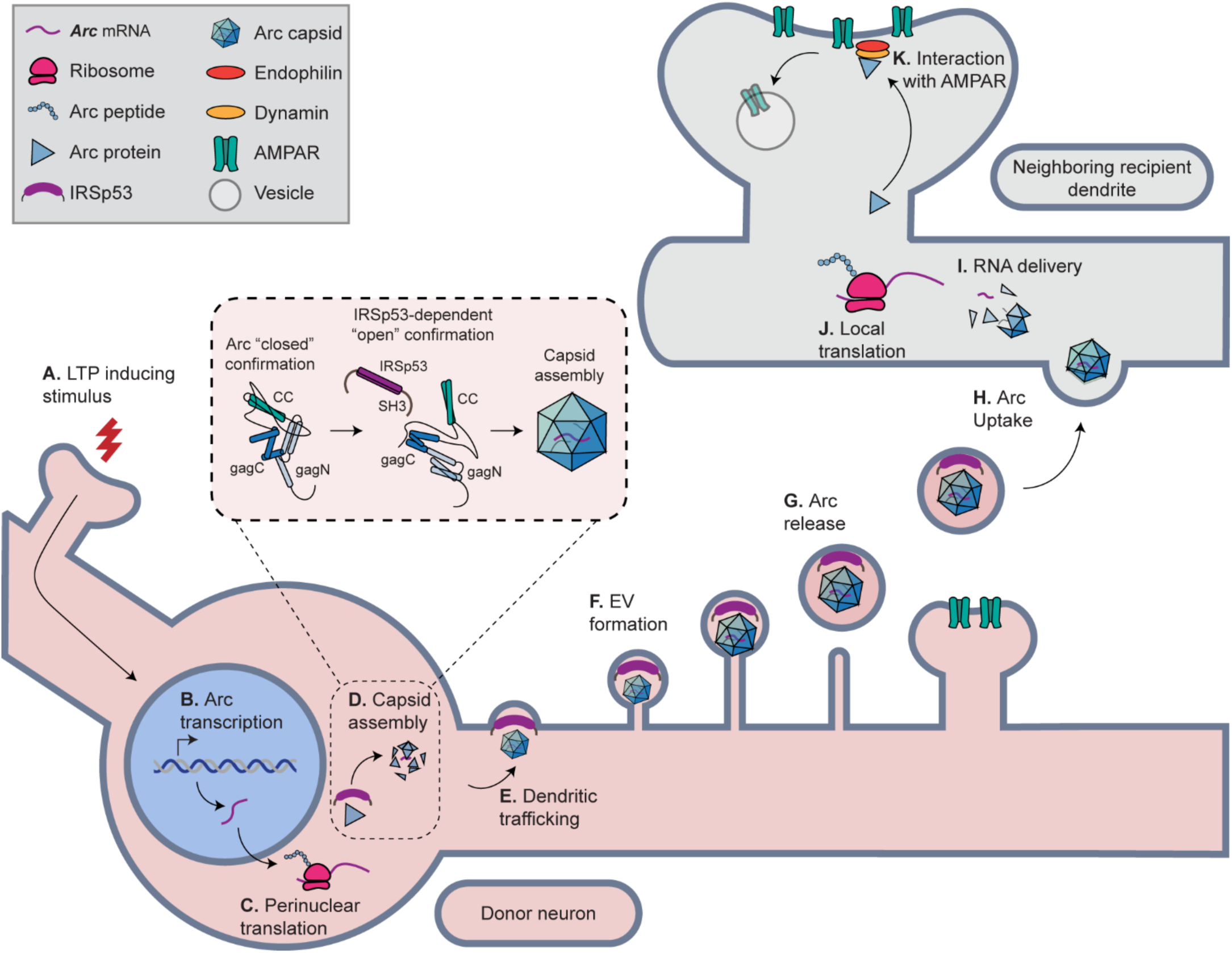
Model of Arc intercellular synaptic plasticity. **A**. LTP induction results in downstream signaling cascades that includes: **B**. activity-dependent Arc transcription. **C**. *Arc* mRNA is transported to the perinuclear cytoplasm where it is translated into protein and interacts with IRSp53. **D**. IRSp53 binding facilitates Arc capsid assembly by switching Arc into an “open” conformation for capsid-prone Gag interactions **E**. Arc capsids are trafficked to dendrites in an IRSp53-dependent manner. **F**. In dendrites, Arc and IRSp53 interact with the membrane at actin-rich extra-synaptic microdomains that dynamically form filopodia. **G**. Arc capsids are released from filopodia. **H**. Arc EVs are taken up by nearby neighboring dendrites via endocytosis. **I**. Arc capsids are disassembled in the endosome and deliver *Arc* mRNA into the cytoplasm. **J**. *Arc* mRNA is then locally translated into protein that (**K**) accelerates the endocytosis of AMPARs from the synapse, resulting in selective weakening of synapses.

## Discussion

EV intercellular signaling plays a role in diverse biological processes, ranging from plant defenses against microbes^42^ to the spread of pathology in neurodegenerative disorders^43^. However, the function of neuronal EVs remains poorly described, and their role in the nervous system is unclear^44^. This is partly due to a lack of mechanistic understanding of the biogenesis and cargo of select EVs. The unexpected discovery that Arc forms virus-like capsids that are released from neurons in EVs^13^ implied that there must be an Arc-dependent intercellular function in the brain. Here, we find that IRSp53 facilitates Arc capsid assembly and Arc EV biogenesis. LTP stimuli induce the formation of Arc-IRSp53 complexes that traffic anterograde into dendrites, where release occurs at actin-rich filopodia. Arc transfer occurs in neighboring dendrites, resulting in a loss of surface AMPA receptors. Arc EVs are sufficient to induce loss of surface AMPARs in recipient neurons, which requires delivery and translation of *Arc* mRNA. Together, these results suggest that Arc mediates a novel form of intercellular synaptic plasticity through LTP-induced IRSp53-dependent release of Arc EVs (Figure S8). We also reveal a central role for IRSp53 in regulating Arc capsid assembly, trafficking, and release into EVs from the plasma membrane – identifying a new mechanism for neuronal EV biogenesis.

### IRSp53 facilitates Arc capsid assembly through a conformational switch

Retroviruses use scaffolding molecules, such as inositol hexakisphosphate (IP_6_) or nucleocapsid proteins, to provide structural support during capsid assembly^45,46^. Mammalian Arc has structural homology to HIV capsids, and *Drosophila* Arc proteins assemble retrovirus-like icosahedral capsids^47^. Here, we find that IRSp53 protein directly facilitates the assembly of Arc capsids, likely by binding the Arc N-terminal CC region, which switches monomeric Arc into an “open” conformation that allows Gag domain oligomerization. Whether IRSp53 plays a similar function in retrovirus capsid assembly remains to be determined, but we note that this mechanism differs from higher-order interactions between pentamers and hexamers in HIV1 capsid assembly^48^. We propose an autoinhibitory mechanism of Arc monomers, through a large-scale structural rearrangement, which enables separation of the Arc N-terminus from the Gag domain. In contrast, the HIV capsid mechanism involves “refolding” of the regulatory TVGG motif, which enables a ratcheting mechanism and reshaping of the NTD-NTD/NTD-CTD interfaces^48^. This TVGG motif is 150-155 residues in mouse Arc (Q8BKX1) and is not involved in the regulatory mechanism we propose here. Our data also shows that the role of IRSp53 in EV release can be dissociated from its role in Arc capsid assembly, as the IRSp53^I403P^ mutant does not enhance Arc release, presumably due to a lack of downstream actin regulation through SH3 domain interactions^37^, but can still bind Arc and facilitate Arc capsid assembly. Further support for this model comes from the Arc^Δ20–40^ N-terminal deletion that removes the CC inhibitory domain, allowing Arc to readily form capsids without IRSp53 binding.

### IRSp53 is required for LTP-induced Arc anterograde trafficking and EV release from dendrites

EVs have been implicated in synaptic function through intercellular transfer of synaptobrevin^49^ and regulation of Wnt signaling through the protein proline-rich 7^50^. Previous studies showed that AMPARs may also be released in EVs^51^, and neuron-derived exosomes may also be critical for neural circuit development^52^. Here, we reveal a new EV biogenesis mechanism regulated by IRSp53. IRSp53 is an I-BAR scaffolding protein highly expressed in neurons that regulates dendritic spine formation and NMDA receptor function^20,53,54^. Recent studies have implicated IRSp53 in EV release in non-neuronal cells via direct budding from the plasma membrane^22–24^. Our data reveals a neuronal activity-dependent EV biogenesis pathway – LTP induction results in the formation of Arc-IRSp53 complexes, which traffic anterograde into dendrites where release occurs from dynamic filopodial and actin-rich protrusions. However, we do not definitively know where and when the fully assembled Arc is released, as our live imaging experiments do not have this spatial resolution.

Interestingly, loss of IRSp53 prevented activity-dependent Arc trafficking into dendrites. This may be due to a requirement for the trafficking of Arc in a specific biophysical state, indicating that the somatic pool of translated Arc protein is the source of Arc in EVs, rather than locally translated dendritic Arc protein. This spatiotemporal protein trafficking may specify where and when Arc EVs are released in response to neuronal activity and could explain why Arc expression induced by LTP does not immediately result in the removal of AMPARs^55^, as the process of Arc intercellular plasticity takes hours and we predict will be synapse specific.

We note that, similar to retroviruses, Arc release may occur via different routes in different cell-types. For example, Arc release from endosomal-derived exosomes was observed in HEK293 cells that overexpress Arc^56^, although HEK293 cells do not normally express Arc and release was not blocked by ESCRT expression^56^. Our data in neurons and HEK293 cells is not consistent with the canonical multi-vesicular body origins of exosomes, as Arc release is not ESCRT-dependent. In our live imaging experiments, we observed the rapid formation of filopodia that contain Arc and IRSp53 proteins, where an abrupt loss of Arc protein occurred suggestive of a release event. This is consistent with non-neuronal cell studies showing that IRSp53 helps induce EV release from filopodia^22^, similar to another I-BAR protein MIM^57,58^. Together, these data reveal a new EV biogenesis pathway in neurons that is more similar to direct “budding” of retroviral capsids from surface plasma membranes than classic exosome release from multi-vesicular bodies.

### LTP and LTD differentially regulate Arc function

Arc transcription is clearly induced by LTP stimuli^9,59^. However, there is conflicting evidence for the role of Arc in the expression of LTP itself. LTP is intact in the hippocampus of Arc KO mice, but Arc-specific antisense oligonucleotides cause deficits in dentate gyrus LTP^2,10,60,61^. Arc’s role in LTD is clearer, with well-defined molecular mechanisms implicating Arc in the endocytosis of AMPARs^4,6,41,62^. Here, we resolve some of these discrepancies by showing that LTP, rather than LTD stimuli, preferentially causes the release of Arc. This switch in Arc function is primarily mediated by IRSp53, which catalyzes Arc capsid assembly and trafficking into dendrites in response to LTP but not LTD. This is further supported by the lack of effects on Arc release after LTD, which is known to induce local Arc translation in dendrites that results in AMPAR endocytosis at synapses^63,64^. We posit that synaptic Arc protein regulates AMPAR trafficking in a non-capsid form, as post-synaptic density protein-protein interactions may prevent higher-order Arc oligomerization^65,66^.

### Intercellular synaptic plasticity and memory

Arc is critical for memory consolidation^2,61^ and experience-dependent plasticity^67,68^. Recent studies have shown that Arc transcription can self-regulate cycles of transcription, leading to waves of expression in the same neuron^69^. Thus, Arc synthesis may be sustained in neurons active during memory encoding. Our findings suggest that memory formation may involve intercellular signaling that requires non-cell autonomous regulation of synaptic function – a new element in the mechanisms of memory formation. Recent studies also suggest that intercellular communication can occur through dendritic nanotubes^70^, highlighting the growing mechanisms of cell-to-cell communication in the brain. We predict that the spatial and temporal aspects of Arc release will be critical for regulating intercellular synaptic plasticity. These slow processes take hours from Arc transcription to Arc EV uptake, local translation, and AMPAR endocytosis – on par with the temporal evolution of the molecular consolidation of memory^71^. We observed a tight spatial constraint on Arc’s effect on synapses, even in primary cultured neurons – Arc transfer only occurs at neighboring dendrites. However, the precise specificity of effects on synapses *in vivo* will need to be determined.

We speculate that Arc facilitates a novel form of circuit-level synaptic plasticity that may help consolidate memory by increasing the “signal-to-noise” of neurons in a memory “engram” circuit by weakening synapses neighboring non-engram neurons. This experience-dependent acceleration of synapse turnover may also contribute to representational drift and memory updating^72^. This is similar to the hypothesis that Arc mediates heterosynaptic LTD on the same neuron by selectively removing AMPARs from inactive synapses^73,74^, but we hypothesize that Arc also weakens synapses on non-connected inactive synapses spatially near the active neuron.

## Conclusion

Our data reveals a new EV-dependent intercellular signaling pathway regulated by IRSp53 and Arc. Arc EVs induce loss of surface AMPARs in recipient neurons to mediate an intercellular form of synaptic plasticity. Our findings resolve previous studies showing that LTP induces Arc expression despite not playing a critical role in LTP itself. Our studies implicate non-autonomous intercellular synaptic plasticity in memory consolidation. Future work will test these novel hypotheses and the precise function of Arc EVs *in vivo*.

Our observations lead to a number of open questions. The dependency of *Arc* mRNA transfer for the functional effects on AMPARs suggests that once Arc EVs are taken up, *Arc* mRNA must be delivered and released into the cytoplasm, where it is translated. The precise molecular mechanisms of Arc capsid disassembly and mRNA release from endosomes remain to be elucidated. The specificity of Arc EV uptake *in vivo* also remains to be determined; we hypothesize that unique receptors engage in Arc-EV uptake and delivery that will further constrain the spatial specificity of Arc EV function within a circuit. IRSp53 has been implicated in psychiatric disorders such as autism spectrum disorder (ASD), schizophrenia, and attention-deficit/hyperactivity disorder (ADHD)^20,75,76^. Future studies will determine whether IRSp53-dependent EV biogenesis is disrupted in these disorders. Furthermore, Arc EVs also play a role in the intercellular transmission of tau^16^ and, thus, the IRSp53-Arc interaction may be targeted for potential therapeutic treatments of these brain diseases.

## Acknowledgements

We thank Dr. Hiroki Okuno and Dr. Haruhiko Bito (Tokyo University) for gifting the Arc synaptic activity-responsive element and minimal promoter plasmids. We thank Dr. Richard Huganir (Johns Hopkins Medical School) for providing the N-terminal GluA1 antibodies. We thank Dr. Eunjoon Kim (Korea Advanced Institute of Science and Technology) for gifting us with the *IRSp53^flx/flx^* mice. The authors would like to acknowledge David Rademacher and the Imaging Core at the Stritch School of Medicine for assistance in generating electron micrographs of Hibit-tagged Arc capsids. We acknowledge the electron microscopy and cell imaging cores at the University of Utah for use of equipment (JEOL-JEM TEM, Leica Spinning Disk microscope, and the Zeiss 880 with Airyscan module) and thank David Belnap and Xiang Wang for their technical assistance. We thank all members of the Shepherd Lab, past and present, for technical support and help.

E.M.C, T.G., and J.D.S were supported by a NIH Director’s Office Transformative Research Award (R01 NS115716). K.S. was supported by a NIH F99/K00 award (F99 NS139545). M.F. was supported by AIRC IG 26229. J.D.S was supported by the Chan-Zuckerberg Initiative Ben Barres Early Acceleration Award, the Jon M. Huntsman Presidential Endowed Chair fund, and NIH R01MH112766.

## Author Contributions

K.S performed protein purification, electron microscopy, live-cell imaging, Airyscan imaging, molecular biology, EV isolation, Arc uptake/transfer assays, immunocytochemistry, image analysis, and confocal microscopy. A.R performed the *in vivo* and *in vitro* immunoprecipitation assays, Arc-Hibit assays in HEK293T cells and primary cultured neurons, Arc transfer assays, immunocytochemistry, molecular biology, and confocal microscopy. E.D.H performed the proximity ligation assay, electrophysiology, and molecular biology. S.K conducted the capsid binding assay and the U87 Arc release assay. J.E performed the *IRSp53^flx/flx^*primary neuron EV collections and biochemistry. T.K performed the equilibrium density gradient isolation and nanoparticle tracking analysis on Arc-containing EVs. M.T conducted the IRSp53 GST-pulldown. A. W. performed the characterization of ESARE-Arc and CMV-Arc expression. B.F performed all-atom molecular dynamics to generate the model of Arc and IRSp53, and M.F. conceived the Arc-IRSp53 biophysical model. A. W. and T.K. performed Banker-style co-culture experiments. T.K, T.S, and A.S generated the mouse and rat primary cultured neurons. K.L designed and cloned the Arc-RaPTR reporter and live-cell imaging of Arc protein and mRNA. A.D performed the Arc and IRSp53 colocalization analysis. K.S, A.R, E.D, M.H, J.E, S.K, T.K, K.L, A.D, M.F, T.G E.C, and J.D.S designed, analyzed, and interpreted experiments. K.S, A.R, and J.D.S wrote the manuscript; all authors discussed results. K.S, T.G, E.C, and J.D obtained funding for the project.

## Methods

### Plasmids

Arc-RaPTR (RNA and Protein Translation Reporter): The Arc synaptic activity-responsive element (SARE) and minimal promoter plasmids were obtained from Hiroki Okuno and Haruhiko Bito (University of Tokyo). This plasmid was PCR amplified with Mfe1 and Not1 and inserted into the pUB_smHA_KDM5B_MS2 (Addgene plasmid # 81085) lentiviral vector^77^ replacing the UbC promoter. The 5’ end of this construct contains 10 copies of the HA epitope while the 3’ end contains 24 MS2 RNA stem-loops. The Arc ORF was subcloned from pRK5-Arc and the insert was PCR amplified with AsiS1 and Nhe1 and ligated into the pUB_smHA_KDM5B_MS2 backbone.

Arc deletion mutants were generated from pRK5 expression plasmids containing WT rat Arc. Pairs of overlapping primers with homology arms were designed for each deletion, targeting nucleic acid sequences adjacent to the deletion region. Overlap extension PCR reactions were performed using these primers to amplify the full plasmid Arc ORF and backbones minus the targeted deletion sequence to generate an Arc deletion mutant plasmid.

The short-hairpin RNA targeting Arc (TRCN0000108909) was developed by the Broad Institute and purchased from The RNAi Consortium (TRC). The shRNA sequence, AGTCAGTTGAGGCTCAGCAAT, was packaged into the pLKO.1 lentiviral vector. The short-hairpin RNAs targeting IRSP53 (TRCN0000079054, TRCN0000079056) were also purchased from the TRC. The shRNA sequences are, CGGAAGTGACAGATTGCATAT and CTGAAGAAATACCAAACGGAA, and the scrambled shRNA sequence is GCACGTAAAGAAGAACCTAAAC. IRSp53 shRNAs were cloned into a pLenti backbone by Gibson assembly. Downstream of the shRNAs an hSyn promoter and mScarlet-I3 fluorophore were added via Gibson assembly.

The ORF of mouse IRSp53 (transcript variant 1, NM_001037755) was amplified by PCR with XhoI and BamHI and ligated into the pLVX lentivirus backbone with or without mRuby. IRSp53^I403P^ gene block was generated by IDT and amplified by PCR with XhoI and BamHI. The insert was then ligated into a pLVX backbone. Rat Arc and Arc^Δ20–40^ was amplified by PCR using XhoI and XbaI and ligated into a C-terminal pLVX-HiBiT backbone obtained from the Arc-HiBiT plasmid. pLVX-eGFP-Cre (plasmid #86805) and pLVX-GFP (plasmid #17448) were purchased from Addgene. IRSp53 I-BAR domain was amplified from the pLVX-IRSp53 plasmid and ligated into a pRK5-myc backbone using SalI and NotI.

Protein purification was carried out using the pGEX-6p1 (GE Healthcare, Little Chalfont, UK) bacterial expression vector. The ORF of mouse IRSp53, IRSp53^I403P^, and Arc^Δ20–40^ were amplified by PCR from pRK5 vectors with BamH1 and Not1 and ligated into the pGEX-6p1 vector.

### Antibodies

Western blots – Primary antibodies: Arc (1:200; mouse monoclonal, Santa Cruz; 1:1000; rabbit polyclonal, Synaptic Systems), Alix (1:500; rabbit polyclonal, custom antibody provided by the Dr. Wesley Sundquist), IRSp53 (1:1000, mouse monoclonal, Abcepta; 1:1000, rabbit polyclonal, Proteintech), PSD-95 (1:2000; mouse monoclonal, US Davis/NIH NeuroMab), GFP (1:1000; mouse monoclonal, Millipore Sigma), myc (1:1000, Sigma Aldrich). All secondary antibodies were used at a dilution of 1:10,000 (HRP-conjugated goat anti-rabbit, goat anti-mouse, goat anti-chicken, Jackson ImmunoResearch).

Immunocytochemistry *–* Primary antibodies: Arc (1:1000 anti-guinea pig, Synaptic Systems, 1:1000 anti-rabbit, Synaptic Systems, Goettingen, Germany; 1:1000 anti-rabbit, Arc custom antibody, generated by Thermo Fisher); IRSp53 (1:1000 anti-rabbit, Proteintech, Rosemont, IL), MAP2 (1:2000 anti-chicken, ab5392; Abcam), N-terminal GluA1 (1:500 anti-mouse, custom antibody provided by Dr. Richard Huganir, Johns Hopkins Medical School). Secondary antibodies: Alexa Fluor 405, 488, 555, or 647 for the appropriate animal host (1:1000 Thermofisher Scientific).

### Cell Lines

HEK293T cells were purchased from ATCC (#CRL-11268). Cells were maintained at 37°C/5% CO_2_ in DMEM media supplemented with 10% fetal bovine serum (FBS) (Thermo Fisher). For transfections, HEK cells were seeded at 40% confluency to either 6-well plates or 10cm dishes. HEK cells were transfected using Polyethylenimine (PEI) (1mg/mL) (Polysciences, cat# 23966-2). 3µg plasmid DNA (6-well dishes) or 8µg DNA (10 cm dishes) was diluted in OptiMEM whose volume was 10% of the final volume in culture dish. DNA and PEI incubated for 20 min at room temperature (RT) and added to cells to incubate overnight at 37°C/5% CO_2_. A full media change was performed with DMEM supplemented without FBS at 24 hours and media was collected 48 hours post-transfection.

### Lentivirus

HEK293T cells (cultured in DMEM + 10% FBS; no antibiotics) at 80% confluency in 10cm dishes were co-transfected with 3µg each of psPax2, pHEF-VSV-G, and pLVX-GFP/eGFP-Cre/pLVX-Arc-HiBiT/pLVX-mIRSp53/pLVX-mIRSp53^1403P^/Arc^Δ20–40^ -HiBiT or pLKO.1 Arc-shRNA. Plasmid DNA was diluted in OptiMEM and incubated with PEI for 20 min. at RT. Cells underwent a full media change 24 hours post-transfection without FBS and media was collected 48 hours post-transfection. Media was centrifuged at 1000*xg* for 10 min. at 4°C, HEPES was added to a final concentration of 20µM, and then filtered through a 0.45 micron low protein binding filter. Media was then centrifuged at 5000x*g* overnight at 4°C and the viral pellet was resuspended in 1X phosphate buffered saline (PBS). Lentiviral particles were stored at −80°C until used for experiments.

### Mouse models

Wild-type and Arc KO mice^67,78^ used in these studies were littermates from heterozygous crosses. IRSp53^flx/flx^ mice (a gift from Dr. Eunjoon Kim, Korea Advanced Institute of Science and Technology) were used from homozygous crosses. Both male and female mice were used. Mice were group-housed with littermates of the same sex after weaning (2-5 mice/cage), on a 12:12 hr. day:night cycle, with food and water *ad libitum*.

### Rat models

Taconic Biosciences/Cyagen generated the Arc (GenBank accession number: NM_019361.1; Ensembl: ENSRNOG00000043465) KO rats in a Long Evans background by CRISPR/Cas9. gRNAs were created by i*n vitro* transcription (gRNA1: AAGACCTAGACCCATGTATCTGG, gRNA2: AGGCACAGGAACCCCGCCTATGG) and co-injected into fertilized eggs for Arc KO production. Arc KO rats were confirmed by PCR (F1: 5’-GATAGGTTAGCTCTAAGAGAGGCAG-3’, R1: 5’-TTGTGTACTCTTGGTAATTCACCTCT-3’) with WT allele = 5475 bp and mutant allele = 806 bp. Long Evans Arc KO rats used in this study were from heterozygous crosses. Homozygous pairs were then used for generating primary neuronal cultures. Rats were group-housed with littermates of the same sex after weaning (2-4 rats/cage), on a 12:12 hr. day:night cycle, with food and water *ad libitum*.

### Cultured Primary Neurons

Primary neurons were prepared from E18 mouse/rat cortices and hippocampi as previously described^8^. Tissue was dissociated using 0.01% DNase (Sigma Aldrich) and 0.06*7*% papain (Worthington Biochemicals) prior to trituration through glass pipettes to obtain a single-cell suspension. Hippocampal cells were then plated at 9×10^4^ cells/mL in Neurobasal medium (Thermo Fisher) supplemented with 5% FBS (Thermo Fisher), 2% GlutaMAX (Thermo Fisher), 2% SM1 (Thermo Fisher), 1% penicillin and streptomycin (Thermo Fisher) on coverslips (No. 1, Bioscience Tools) coated overnight with 0.2 mg/mL poly-L-lysine (Sigma-Aldrich) in 200mM borate buffer (pH 8.5). Cortical cells were plated at 4×10^6^ in previously described Neurobasal medium on 10 cm^2^ dishes (Corning) prepped with poly-L-lysine. Neurons were grown at 37°C/5% CO2 and fed via half-media exchange every 3rd day with astrocyte-conditioned BrainPhys media (StemCell Technologies) supplemented with 1% FBS, 2% SM1 (StemCell Technologies), 500 µM L-Glutamine, and 1% penicillin/streptomycin with DIV5 feeding containing 5µM β-D-arabinofuranoside (Sigma-Aldrich) to limit overgrowth of glial cells. Neurons were grown for 14-16 days *in vitro* (DIV) prior to experiments.

### Neuron Transfection

Neurons were transfected after DIV14-15 using lipofectamine 2000 (Thermo Fisher) at a 3:1 ratio when complexed with plasmid DNA. Neurons were transfected over the course of 1 hr. at 37°C in pH 7.2 Minimum Essential Media (Thermo Fisher) supplemented with 2% GlutaMAX, 2% SM1, 15 mM HEPES (Thermo Fisher), 1 mM Sodium Pyruvate (Thermo Fisher), and 33mM Glucose without CO_2_. After transfection, the neurons were given 12 hours in growth media at 37°C/5% CO_2_ to allow sufficient recovery and expression of the plasmid prior to fixation in 4% formaldehyde (Thermo Fisher) / 4% sucrose (VWR) in 1X PBS for 15 min. at RT. After fixation, neurons were washed with PBS and mounted in Fluoromount G Mounting Medium with or without DAPI (Thermo Fisher) for ICC.

### Immunocytochemistry (ICC)

Neurons were fixed by transferring and inverting coverslips to a prepared 12-well plate with 1ml/well of 4% sucrose/4% formaldehyde (ThermoFisher Scientific) in 1X PBS and incubated at RT for 15 minutes with rocking. Neurons were washed 3 x 5 minutes with 1X PBS and stored at 4C for several days. All coverslips were visualized under a light microscope to assess cell health, and the best two from each condition were selected for ICC. Neurons were permeabilized for 10 minutes with 0.2% Triton X-100 (Amresco, Solon, OH) in 1X PBS, and blocked in 10mg/ml bovine serum albumin (BSA) in 1X PBS for 30 minutes at RT. Neurons were incubated in primary antibodies in block for 1 hour at RT, washed 3 x 5 min. in 1X PBS, and incubated in secondary antibodies (donkey anti-chicken Alexa Fluor 488, donkey anti-rabbit Alexa Fluor 647 both 1:2000) diluted in block for 1 hour at RT. Neurons on coverslips were mounted on glass slides in Fluoromount with Dapi (ThermoFisher Scientific) and dried overnight at RT.

### Confocal Imaging and Analysis

Five 6um z-stack (0.5um step) images were taken per experimental group on a Nikon 60X oil objective, with imaging settings set to a no-primary control. Maximum Intensity Projections were generated in ImageJ. Custom ImageJ code was used to identify the nucleus based on the Dapi signal and expand the radius to create an ROI of the entire neuronal soma. Within the soma ROI, the Raw Integrated Density of the Arc channel was measured and divided by the area of the MAP2 mask. The soma ROIs were then subtracted from the image so that only dendrites remained. Two 15×5 micron ROIs were drawn on primary or secondary dendrites of each neuron, 3-5 neurons per image. The Raw Integrated Density of the Arc channel was measured within these ROIs for all conditions, divided by the area of the MAP2 mask within that ROI, and averaged for each neuron. The values from the KO coverslip / KO 10cm dish condition, which represent background signal from the primary antibody, were subtracted from the plotted values for the experimental conditions.

### Extracellular Vesicle Purification

EVs for synaptic plasticity assays were purified from culture media of DIV15 mouse primary cortical neurons. Media was collected and spun sequentially at 2,000*xg* and 20,000*xg* to remove any cell debris and apoptotic bodies. The clarified media was then concentrated to 0.5 mL using 100kDa Vivaspin protein concentrator spin columns (Cytvia). Concentrated EVs were further fractionated using qEV Gen 2 IZON size exclusion columns (IZON). Void fractions were discarded (3mL) and early fractions that contained EV markers and Arc (fractions 1-4) were used for EV uptake assays at a concentration of 50µg, as determined by nanodrop.

EVs from IRSp53^flx/flx^ mouse cultures were collected from media obtained from five 10cm biochemistry dishes and combined into a single EV preparation isolated by ultracentrifugation over a sucrose cushion. Differential Centrifugation was used to isolate the smallest components released from cells. Low-speed spin (2000*xg* for 10 min.) is performed to remove the largest components in cell culture media followed by a higher-speed spin (20,000*xg* for 20 min.). This was followed by a 134,000*xg* for 70 mins. and then a 33,000*xg* spin on a 20% sucrose cushion for 16 hours to pellet EVs in the 100-120nm size range.

### HiBiT assays

Arc-HibiT samples from HEK293T cells were collected 48 hours post transfection and neurons 6 days post transduction. Conditioned media from HEK293T cells or neurons, plated in 6-well dishes were centrifuged at 1000*xg* for 10 minutes and suspended in 5x Passive Lysis buffer (Promega). Each data point represents an individual well. Arc-HiBiT signal was identified by complementation with LgBiT (equal volumes) and measurement of Nanoluc-mediated luciferase (Nano-Glo Luciferase Assay, Promega, 565nm) on a microplate reader (SpectraMax iD3). Total Arc-HiBiT in the cell lysate is calculated by dividing the luminescence value in the cell lysate by the percentage of the sample used from the total sample collected. Total Arc-HiBiT in the media is calculated by dividing the media luminescence value by the percentage of the media sample used from the total sample. The percentage of Arc release is calculated by dividing the total luminescence value in the media by the sum of the total luminescence in the cell lysate and the total luminescence in the media.

Arc EV HiBiT assay: HEK293T cells were seeded in 10 cm cell culture dishes at ∼80% confluency. The cells were transfected with Arc-HiBiT plasmid and either IRSp53 or empty vector plasmids using LipoD293 reagent (SignaGen) according to the manufacturers protocol. 48 hours post transfection, conditioned media from Arc-HiBiT expressing HEK293T cells was harvested and clarified by sequential 300*xg* and 3000*xg* spins. Clarified media was overlayed onto 20-50% w/w sucrose gradients (prepared in serum free DMEM). The gradients were centrifuged in an SW41 rotor at 200,000*xg* for 1hr. Fractions were collected from air-gradient interfaces and complemented using LgBiT (Promega, 1:200 dilution) and nanoluc substrate (Promega, 1:200) diluted in 1X passive lysis buffer (Promega). Luminescence of the fractions was measured with a luminometer (Promega, GloMax Explorer). EV-associated Arc-HiBiT was identified in gradient fractions 15-17 as these fractions have a density consistent with the known densities of EVs (1.10-1.15g/mL).

### EV nanoparticle tracking analysis

HEK293T cells were seeded in 10 cm cell culture dishes at ∼80% confluency. The cells were transfected with Arc-HiBiT plasmid and either IRSp53 or empty vector plasmids using LipoD293 reagent (SignaGen) according to manufacturer protocol. 48 hours post transfection, conditioned media from Arc-expressing HEK293T cells was harvested and clarified by sequential 300*xg* and 3000*xg* spins. The clarified supernatant was applied to 100K Amicon ultrafilters and concentrated ∼20-fold. The concentrated media was loaded on IZON qEV original 35 nm size exclusion columns according to manufacturer protocols. Fractions were collected using the IZON automated fraction collector. Fractions were complemented using LgBiT (Promega, 1:200 dilution) and nanoluc substrate (Promega, 1:200) diluted in 1X passive lysis buffer (Promega). Luminescence of the fractions was measured with a luminometer (Promega, GloMax Explorer). EV associated peak fractions were 2-fold serially diluted in PBS and analyzed by nanoparticle tracking analysis (Nanosight NS300, Malvern) to calculate average diameter and concentration of particles. Particle tracks (n = 3) are depicted as averages (line traces) with standard errors of the means depicted as shaded areas.

### Western blots

Western blot samples were mixed with 4X Laemmli buffer (40% glycerol, 250 mM Tris, 4% SDS, 50 mM DTT, pH 6.8) and heated at 70°C for 5 min. SDS-PAGE gel electrophoresis was used to separate protein samples. Separated protein samples were transferred to a nitrocellulose membrane (GE Healthcare). Following transfer, membranes were briefly stained with Pierce™ Reversible Total Protein Stain and then destained for total protein imaging. Membranes were blocked in 5% milk + 1X Tris-buffered saline (TBS; 10x: 152.3 mM Tris-HCl, 46.2 mM Tris-base, 1.5 NaCl mM, pH 7.6) for 30 min. at RT. Membranes were then incubated with in primary antibody in 1X TBS overnight at 4°C. Membranes were washed 3 x 10 min. in 1X TBS, then incubated with in an HRP-conjugated secondary antibody (Jackson ImmunoResearch) in block for 1 hour. at RT. After 3 x 10 min. washes in 1X TBS, a chemiluminescence kit (Bio-Rad, Hercules, CA) was used to detect the protein bands. Membranes were imaged on an Amersham ImageQuant 800 gel dock (Cytiva). Blots were analyzed using ImageJ/FIJI (National Institutes of Health, Bethesda, MD).

### Immunoprecipitation

Mouse forebrain tissue was dissected out and homogenized in immunoprecipitation (IP) lysis buffer (150 mM NaCl, 50 mM Tris, 3% Triton X-100, pH 7.4), with protease inhibitor cocktail added fresh (Roche). Homogenates were pelleted at 200*xg* for 5 min. at 4°C to remove tissue debris. Supernatants were removed and rotated for 10 min. at 4°C before being pelleted at 10,000*xg* for 10 min. at 4°C to remove insoluble material. Cleared supernatants were removed and a small aliquot was taken as the input, while the remainder was used for the co-IP. Supernatants were immunoprecipitated either with Arc antibody (rabbit polyclonal, custom-made; ThermoFisher) or normal rabbit IgG (Santa Cruz Biotechnology, Santa Cruz, CA) at 2µg/mL overnight at 4°C while rotating. Following antibody incubation, a 10% volume of washed Protein A/G magnetic bead slurry (ThermoFisher Scientific) was added to the antibody/lysate mixture and incubated for 1 hour at 4°C rotating. Bead-antibody mixture was washed 3 times with IP lysis buffer and beads were resuspended in 200µL IP buffer and then 4X Laemmli buffer was added. Protein was separated from magnetic beads at 70°C for 5 min. and then the beads were isolated using a magnetic bead stand. Input (10% lysate volume) and each of the IgG and Arc antibody elution’s were separated by SDS-PAGE on a 10% acrylamide gel and immunoblotted as described above.

### Protein Purification

GST-tagged proteins were purified from *Escherichia coli* Rosetta 2 BL21 competent cells as previously described^13^. Starter bacteria cultures were grown overnight at 37°C in LB supplemented with ampicillin and chloramphenicol. Large-scale 500 mL cultures were inoculated with stater cultures in ZY auto-induction media. Large-scale cultures were grown to OD_600_ of 0.6-0.8 at 37°C at 160rpm then shifted to 18°C at 180rpm for 16-20 hours. Cultures were pelleted at 4000x*g* for 15 min at 4°C. Pellets were resuspended in 25 mL of lysis buffer (500 mM NaCl, 50 mM Tris, 5% glycerol, 1 mM DTT, pH 8.0) and flash frozen in liquid nitrogen. Frozen pellets were thawed in 37°C and brought to a final volume of 1g pellet:10 mL lysis buffer, supplemented with DNAse, lysozyme, and complete protease inhibitor cocktail (Roche). Resuspended lysates were sonicated for 6 x 45s pulses at 90% duty cycle and pelleted for 75 min. at 21,000x*g*. Cleared supernatants were incubated with pre equilibrated GST Sepharose 4B affinity resin in a gravity flow column overnight at 4°C. GST-bound protein was washed twice with 20 resin bed volumes of lysis buffer and re-equilibrated in TBS (150 mM NaCl, 50 mM Tris, 1 mM EDTA, 1mM DTT, pH 7.2) at RT and cleaved from GST resin overnight at 4°C with PreScission Protease (GE Healthcare). GST was affinity purified as described above and eluted using 15 mM reduced L-gluathione, 10 mM Tris, pH 7.4 at RT.

### GST-pulldown assays

Proteins were purified with the GST tag intact or were cleaved using PreScission protease. GST-tagged proteins were incubated with specific cleaved proteins overnight at 4°C. The following day, a column was washed several times with TBS, and the incubated proteins were added to the column. PreScission protease (L-reduced glutathione (20mM, pH 8.0)) was added to the column and incubated with the proteins for 20 min. The cleaved protein complexes were eluted from the column and prepared for SDS-PAGE and western blot.

### Arc Capsid Assembly Assay

GST cleaved purified Arc and IRSp53 protein elutes were concentrated in Vivaspin 30kDa MWCO ultrafiltration spin columns (Cytvia) to 2.5 mL and protein concentration was measured by nanodrop. Equal concentrations of Arc, IRSp53, or Arc and IRSp53 incubated in a 1:1 ratio for 2 hours in TBS and were then run on a Superdex 200pg HiLoad 16/600 size exclusion column in TBS. Peak fractions, determined by chromatogram and Coomassie staining, were pooled and concentrated. Following size exclusion, negative stain EM grids were prepared for each sample at 0.5mg/mL and capsid formation was quantified by manual counting of 15 images/grid.

### Negative Stain Electron Microscopy

For all negative stain EM samples, copper 200-mesh grids (Electron Microscopy Sciences) were glow discharged for 25 s in a vacuum chamber at 30 mA. 3.5µL of sample was applied to the grid for 45s. Grids were washed 2 x 5 s with 30µL water droplets, followed by one wash with 1% uranyl acetate (UA) on parafilm. Excess water/UA was removed from the grid with filter paper, and a final droplet of UA was added to grids for 30s. Excess UA was removed, and grids were air dried for 30s. Imaging was performed using either an FEI T12, FEI Tecnai Spirit microscope at 120 kV or a JEOL-JEM 1400 electron microscope. For each experiment, 15 images were taken of each sample from a stereotyped pattern of grid squares. Arc capsids between 30-50 nm were manually counted using ImageJ/FIJI.

### Capsid Binding Assay

Arc proteins were purified as described above and they were subjected to Superdex 200 column in the presence of 500 mM phosphate buffer. Early fractions were collected, and capsid assembly was confirmed by TEM images. HEK293Ts were transfected using PEI to overexpress IRSp53. The cells were lysed by passive lysis buffer containing a protease inhibitor cocktail. The cell lysate was incubated with *in vitro* assembled Arc capsids for 1 hour at 4°C. The mixture was pelleted by ultracentrifugation at 21,000*xg* for 15 min. The pellets were washed with 500 mM phosphate buffer 3x, and they were resuspended in loading buffer for western blot analysis. The supernatant fractions were collected for further analysis. Pellet and input fractions (10x diluted) are analyzed by SDS-PAGE and immunoblot.

### GluA1 Live Surface-Labeling

Primary hippocampal mouse neurons were surface labeled for GluA1 with a live antibody feeding assay as previously described^7,8^. Living DIV14-16 neurons were washed briefly with cooled (10°C) neuronal incubation media (NIM) (1X MEM, 2% GlutaMax, 10mM HEPES, 1 mM sodium pyruvate, 33 mM glucose, 2% SM1 supplement, pH 7.2, filter sterilize) and then incubated in GluA1-NT mouse monoclonal antibody (1:500) (custom provided by Richard Huganir, clone 4.9D) in NIM at 10°C for 20 min. Neurons were washed in NIM and then fixed in 4% formaldehyde prior to immunocytochemistry.

#### Chemical LTP and LTD

Chemical LTP (cLTP) was induced by incubating neurons in tetrodotoxin (TTX) (1:1000) overnight. The following day, neurons were washed 3 times with glia-conditioned BrainPhys (GCBP) media and then given a 5-minute 200µM glycine pulse in GCBP. Neurons were washed an additional 3 times with GCBP and allowed to recover for 30 minutes (GluA1 surface-labeled neurons) or 2 hours (Arc release). Chemical LTD (cLTD) was induced by washing neurons 3 times with GCBP prior to a 5-minute 50µM DHPG pulse. Neurons were washed 3 times and recovered for 30 min.

### Electrophysiology

DIV 13-14 *Arc* knockout hippocampal neurons were incubated with culture media that contained 50 μg of EVs isolated from WT or *Arc* KO neurons for 2 hours. After 2 hours of incubation, neurons were maintained in EV-free media for 1 hour. Whole-cell patch clamp recordings were then obtained from EV incubated neurons for 1 hour that were constantly perfused with aCSF containing (in mM) 125 NaCl, 2.5 KCl, 26 NaHCO_3_, 1.25 NaH_2_PO_4_, 2 MgSO_4_, 2 CaCl_2_, 10 Glucose, 10 HEPES, (pH 7.4; 300 mOsm) at a temperature of 28-31 °C with a flow rate of 1.5 ml/min. ACSF contained 1 μM TTX and 50 μM PTX to block action potentials and GABAergic synaptic transmission, respectively. Gigaohm seal formation was achieved with patch pipettes (3-6 MΩ) filled with internal solution containing (in mM) 120 CsMeSO_3_, 20 CsCl, 2 MgCl_2_, 10 HEPES, 7 phosphocreatine, 0.2 EGTA, 4 Na_2_-ATP, 0.3 Na_2_-GTP (pH 7.2; 280-290 mOsm). Following seal formation and patch pipette capacitance compensation, whole cell breakthrough was achieved. Cells were then held at −70 mV and access resistance was maintained at ≤; 15 MΩ with a ≤; 20% change in access resistance throughout the recording. Recordings were obtained with a Multiclamp 700B and a Digidata 1440A (Molecular Devices), sampled at 25 kHz, and low pass filtered at 2 kHz. Three minutes of mEPSC events was analyzed with python code that incorporated the MiniML neural network^79^ and Graph Pad Prism 10. Cumulative probability plots of event amplitude and inter event intervals were generated by binning events from each cell into 2 pA and 200 ms intervals, respectively.

### Immunocytochemistry

Neurons were washed once with 4% sucrose/1X PBS at 37°C and then fixed for 15 min. with 4% sucrose/4% formaldehyde (ThermoFisher Scientific) in 1X PBS. Neurons were washed 3 x 5 minutes with 1X PBS, permeabilized for 10 minutes with 0.2% Triton X-100 (Amresco, Solon, OH) in 1X PBS and blocked in 5% bovine serum albumin (BSA) in 1X PBS for 30 minutes at room temperature. Neurons were incubated in primary antibody diluted in block for 1 hour at room temperature, washed 3 x 5 minutes in 1X PBS, and incubated in secondary antibody diluted in block for 1 hour at RT. Neurons on coverslips were mounted on glass slides in Fluoromount (ThermoFisher Scientific) and dried overnight at room temperature.

### Fixed Cell Imaging and Analysis

Coverslips were imaged on a 60x/1.4NA oil objective on a Nikon Eclipse Ti2-E inverted confocal microscope and images were analyzed using ImageJ/FIJI software. Neurons were selected for analysis by looking at MAP2 dendritic staining for cell health. Each experimental group was thresholded based on no primary controls and the brightest immunofluorescent sample was used to determine the image acquisition settings in each experiment. All images from a single experiment were based upon these exact same settings.

Analysis of Arc and IRSp53 protein expression – During image analysis images were thresholded (to minimize background fluorescence and normalize the linear range) to the brightest immunofluorescence in an experiment. This threshold was then applied to all other images. Raw integrated density of three 20µm tertiary dendritic segments/neuron were measured for each image. This value was divided by the area of the dendritic segment, determined by MAP2. The control group in each experiment was set to “1”, and the raw integrated density divided by area of all other groups were normalized to this and displayed ± SEM. The Inferno look-up table (LUT) in ImageJ/FIJI was used to highlight expression changes in the representative images.

Analysis of Arc and IRSp53 colocalization *–* An algorithm was designed using the surface feature function in Imaris v8.4.1 (Bitplane) to generate surfaces around Arc signal and the maximum fluorescence intensity of IRSp53 present within these individual Arc surfaces was determined. The algorithm was applied to all images within the same experiment. Background levels of IRSp53 fluorescent intensity were determined and used to calculate the percentage that colocalized with Arc and the percentage that did not colocalize. The same procedure was performed for all experimental conditions and replicates.

### Live-Cell Imaging and Analysis

Live-cell imaging was performed using a Leica Yokogawa CSU-W1 spinning disk confocal microscope with a Leica Plan-Apochromat 63x/1.4 NA oil objective and iXon Life 888 EMCCD camera. DIV 14-16 primary cultured hippocampal neurons were used for all live-cell imaging experiments and were maintained at 37°C/5% CO_2_ with a stage top incubator. For IRSp53-shRNA experiments, neurons were transfected with 1.5µg of ESARE-mNeon-Arc and 1.5µg of hSyn-mScarlet, IRSp53-shRNA, or IRSp53-scrambled control-shRNA. Transfected neurons were immediately treated with TTX (1:1000) and left overnight at 37°C/5% CO_2._ The following day neurons were imaged for 15 minutes while still in TTX. Neurons were then washed to remove residual TTX and incubated in 200µM glycine for 5 minutes to induce cLTP. Neurons were washed again and incubated in glia-conditioned BrainPhys media for the remaining image acquisition for 2 hours. Image drift was accounted for by resetting image coordinates. For Arc-RaPTR experiments, neurons were transfected at DIV14 with 1µg of Arc-RaPTR, HA-GFP Frankenbody, MS2-miRFP703 coat protein, or in combination with a C-terminally tagged IRSp53-mRuby. Images were acquired as described above.

Live imaging experiments were analyzed using ImageJ/FIJI. Images were run through a custom macro to quantify mNeon-Arc intensity throughout the time course. Arc raw integrated density (RID) was measured within a mask of mScarlet. The RID during the glycine pulse (cLTP) was normalized to the RID of the TTX time frames. To quantify changes in filopodial formation, time-lapse max projection images were generated of dendritic ROIs during TTX and after cLTP induction for all conditions. mScarlet area was measured as a proxy for changes in overall filopodial dynamics. For Arc-RaPTR experiments, images were run through a custom macro to threshold and remove background fluorescence for each individual experiment. Kymographs were generated using the KymographClear/KymographDirect ImageJ plugin and software package. Raw integrated density of Arc within a mRuby-IRSp53 mask was measured during the time course. An average raw integrated density of the 5 minutes immediately after the glycine pulse and the last 5 minutes of the image acquisition was calculated.

### Airyscan Imaging and Analysis

Airyscan imaging was performed on a Zeiss LSM880 confocal microscope with Airyscan module using a Plan-Apochromat 63X/1.3 oil M27 objective lens. Images were taken with both traditional LSM confocal imaging as well as using the Airyscan module. The Airyscan uses a 32-channel gallium arsenide phosphide photomultiplier tube (GaAsP-PMT) detector and increases resolution by approximately 1.7x. Normal confocal images were taken first to ensure proper neuronal health. Airyscan dendritic images were taken from the same dendrites of the traditional LSM images. All Airyscan images were analyzed using ImageJ/FIJI. A custom macro was developed to quantify the amount of colocalized Arc, IRSp53, and Actin or PSD95. A mask of MAP2 was generated in each dendritic ROI and subtracted from the image leaving only extra-dendritic signal remaining. Colocalization was measured as the total overlapping pixel area generated by pixel multiplication between the other channels for each dendritic ROI.

### Proximity Ligation Assay

Neurons underwent cLTP, Basal, or cLTD, and were fixed after 1 hour and underwent Duolink proximity ligation (Sigma). Following fixation, permeabilization, and blocking, primary rat hippocampal neurons were incubated with primary antibodies for Arc and IRSp53 for 1 hour at room temperature. Cells were washed (2x) with wash buffer. Coverslips were then incubated with secondary antibodies for 1 hour at 37°C. These antibodies contain DNA fragments that will ligate when each protein of interest is within 40 nm of the other. Cells were then (2x) washed with wash buffer. Coverslips underwent ligation for 30 minutes at 37°C and were washed (2x) with wash buffer. Cells then underwent rolling circle amplification and probe annealing (Texas Red fluorphor) for 100 minutes at 37°C. The following day, cells underwent immunocytochemistry for MAP2 to label cell bodies and dendrites (see above). Cells were mounted with mounting media containing DAPI and imaged. Max intensity projections of the MAP2, PLA, and DAPI channels were thresholded and binarized. After being binarized, the MAP2 mask was dilated, and the DAPI mask was then subtracted from this MAP2 mask (to remove nonspecific PLA staining in the nucleus). The binarized PLA image was multiplied by the MAP2 mask, resulting in a PLA binarized image that only contains PLA particles that were colocalized with MAP2 signal. Dilate was then applied to this image and the number of particles was quantified and normalized to MAP2 area.

### Computational Structure Modeling

Structure predictions were made using the sequences of mouse Arc (UniProt: Q9WV31) and IRSp53 (UniProt: Q8BKX1). We predicted structures by AlphaFold 3 (AF3)^31^for free Arc; in complex with IRSp53; and Arc^20–40^ segment with IRSp53 SH3 domain (375-438 residues). In addition, we predicted Arc^20–40^ in complex with full IRSp53. For each system, 500 structures were generated using 50 random seeds and 10 diffusion samples.

### Molecular Dynamics Simulations

Molecular dynamics simulations were initiated using the three AF3 structures with highest scores (AF3 score: 0.8×"ipTM"+0.2×pTM+0.5×disorder-100×has_clash)^31^. The structures were placed in a rhombic dodecahedral box, with minimum 10nm separation between the molecules and the simulation cell edge. The structures were solvated in water and 150 mM NaCl. MD simulations were conducted using CHARMM36m^80^ and a variant of the TIP3Pε water model^81^ using Gromacs 2022.5. The initial configurations were energy minimized using steepest descent algorithm with position restraints until the maximum force on an atom was below 1000 kJ mol^-1^ nm^-1^. This was followed by an 1875 ps equilibration with gradual release of the 1000 kJ mol^-1^ position restraints. MD simulations were performed with a time step of 2 fs, distinct velocity-rescaling thermostats^82^ for the protein and solvent with 1 ps time constant, and an isotropic Parrinello-Rahman barostat with 1 bar reference pressure and 5 ps time constant. A real-space pair-interaction cut-off distance of 1.2 nm was applied to both van-der-Waals and electrostatic interactions. The long-range contribution to the electrostatic energy was considered using a particle-mesh Ewald (PME) algorithm. The Arc monomer and the Arc^20–40^ + IRSp53-SH3 systems were simulated for 1 μs, while the full Arc + IRSp53 simulations were run for 0.5 μs. The first 100 ns of each replica was discarded from each trajectory. The MDAnalysis Python package was used to analyze the interatomic distances. The protein-protein contact maps were computed using ConAn^83^. Specificity and proline binding pockets were identified based on sequence similarity to other SH3 domains^84^.

### Statistical analysis

One and two-way ANOVAs (including A Kruskal-Wallis one-way ANOVA) was used to compare between groups with repeated measures (with *post hoc* Sidak’s, Tukey’s and Dunnett’s tests), paired t-tests or two-tailed unpaired t-tests (with *post hoc* Welch’s tests) were performed using GraphPad Prism (GraphPad Software, San Diego, CA). Outliers were identified and removed using Prism. For all statistical analysis significance was set at p < 0.05 and all data shown are representative of at least three biologic replicates.

